# Aster repulsion drives local ordering in an active system

**DOI:** 10.1101/2020.06.04.133579

**Authors:** Jorge de-Carvalho, Sham Tlili, Lars Hufnagel, Timothy E. Saunders, Ivo A. Telley

## Abstract

Biological systems are a form of active matter, which often undergo rapid changes in their material state, *e.g*. liquid to solid transitions. Yet, such systems often also display remarkably ordered structures. It remains an open question as to how local ordering occurs within active systems. Here, we utilise the rapid early development of *Drosophila melanogaster* embryos to uncover the mechanisms driving short-ranged order. During syncytial stage, nuclei synchronously divide (within a single cell defined by the ellipsoidal eggshell) for nine cycles after which most of the nuclei reach the cell cortex. Despite the rapid nuclear division and repositioning, the spatial pattern of nuclei at the cortex is highly regular. Such precision is important for subsequent cellularisation and morphological transformations. We utilise *ex vivo* explants and mutant embryos to reveal that microtubule asters ensure the regular distribution and maintenance of nuclear positions in the embryo. For large networks of nuclei, such as in the embryo, we predict – and experimentally verify – the formation of force chains. The *ex vivo* extracts enabled us to deduce the force potential between single asters. We use this to predict how the nuclear division axis orientation in small *ex vivo* systems depend on aster number. Finally, we demonstrate that, upon nucleus removal from the cortex, microtubule force potentials can reorient subsequent nuclear divisions to minimise the size of pattern defects. Overall, we show that short-ranged microtubule-mediated repulsive interactions between asters can drive ordering within an active system.

## Introduction

Proliferation of the genome is a cornerstone of early development in all animals, generally achieved by cell division. Almost all insects first segregate genome copies into hundreds of nuclei (syncytium) and only at a specific nuclear density transform the single cell into a tissue^1^. The distribution and separation of nuclei in syncytia typically display surprising positional uniformity^2–4^. This uniformity is likely required for precise gene patterning and fate determination^5^. How the embryo controls internuclear distance so robustly has been a decades-long topic of debate^6–12^.

In *Drosophila melanogaster*, early nuclear divisions are meta-synchronous, whereby nuclei gradually fill the inner cellular space until, nine division cycles or ~80 minutes post-fertilisation, 300–400 of them arrive at the cell cortex^13^. The nuclei are subsequently embedded within a two-dimensional topology near the cortex of the ellipsoidal embryo, where they undergo four more rounds of division to generate ~6000 nuclei^14^ prior to cellularisation. Some nuclei do not reach the cell cortex, resulting in the final number after 13 divisions at the cortex being less than 8192 (2^13^). The spatiotemporal synchronisation of nuclear divisions is governed by a reaction-diffusion process emerging from nuclei^4^. Furthermore, global nuclear positioning in the early embryo is crucial for synchronisation^12^.

Microtubule dynamics is critical for nuclear migration to the cortex at nuclear cycle (n.c.) 9 and for their regular distribution upon arrival^15–17^. Nuclei are embedded in a regular matrix that either pulls or pushes them apart, leading to precise internuclear distances. In interphase of the cell cycle, microtubules are organised in radial arrays called ‘asters’, which are nucleated and organised by the centrosome^18^. The centrosome acts as the main microtubule organising centre (MTOC) and promotes polymerisation and focusing of microtubules^19^. Two asters are linked to each nucleus, being functional elements of the bipolar spindle during mitosis. We have shown previously that asters are required for efficient separation of daughter nuclei following chromosome segregation, and we hypothesised asters pulling on daughter nuclei^20^. However, it has remained open whether asters are necessary to maintain the distance to neighbouring nuclei, of which on average six exist for each nucleus^3^. Embryos lacking core centrosomal components do not regularly distribute nuclei and abort development after a few division cycles^21–23^. Conversely, embryos that do not form actin caps and membrane furrowing, which are the precursor of uninuclear cell formation during cycle 14 and are thought to help separate nuclei at the cortex, still show regular nuclear distribution at n.c. 10 and 11 ^8^. The causality and the mode of mechanical separation during and after nuclear division remains an open problem, primarily due to limited visualisation in living samples and the growing mechanical complexity during development.

Here, by exploiting embryonic explants^24^, which reduces complexity, and a cell cycle regulation mutant^25,26^ to uncouple microtubule organisation from nuclear division, we determine how microtubule interactions can spatially organise nuclei. In this system, we uncover the physical principles of separation for simple nuclear arrays, reveal the positional autonomy of asters and derive the microtubule-driven mechanical separation potential. We find that nuclei behave as cargo associated to self-organising microtubule asters which have repulsive properties. These results contrast with uninuclear model systems, where geometry, cortical pulling and hydrodynamic forces appear to drive aster movement and centring^27–31^. Our work reveals the underlying local biophysical interactions that pack and order nuclei within a rapidly changing active system.

## Results

Synchronous nuclear duplication within a constant surface area poses a geometrical challenge (Fig. 1A). Spindle elongation during nuclear division^20^ should cause transiently smaller distances between spindles (and their asters) unless some leave the surface after division. Alternatively, spindles may reorient their division axis to optimise the spacing between nuclei. Thus, we quantified neighbour distances (Fig. 1B), focusing on the distance *d* between centrosomes belonging to nearest neighbour nuclei (‘non-sister’). Importantly, despite organelle duplication, synchronous spindle expansion (defined by *s*, Fig 1B), and apparent collective nuclear movement in a finite space, the distance distribution between neighbouring centrosomes within n.c. 10 did not exhibit noticeable decrease as a compensation (Fig. 1C, black dots). Similarly, or possibly as a result, the nuclear separation distance *D* only mildly decreased (Fig. 1D, black dots) during anaphase and telophase when nuclear duplication and mitotic separation occurs (Fig. 1D, blue dots). In subsequent division cycles, the mean distance between centrosomes gradually decreased while nuclear density doubled. Still, we observed no abrupt decrease or oscillation in the nuclear separation during duplication (Fig. 1E, Suppl. Fig. 1). These measurements suggest that a rigid mechanical connection exists between centrosomes. Thus, we hypothesised the presence of a repulsive mechanism between neighbouring nuclei or asters in the highly viscous, and effectively over-damped cytoplasm.

**Figure 1 –.**
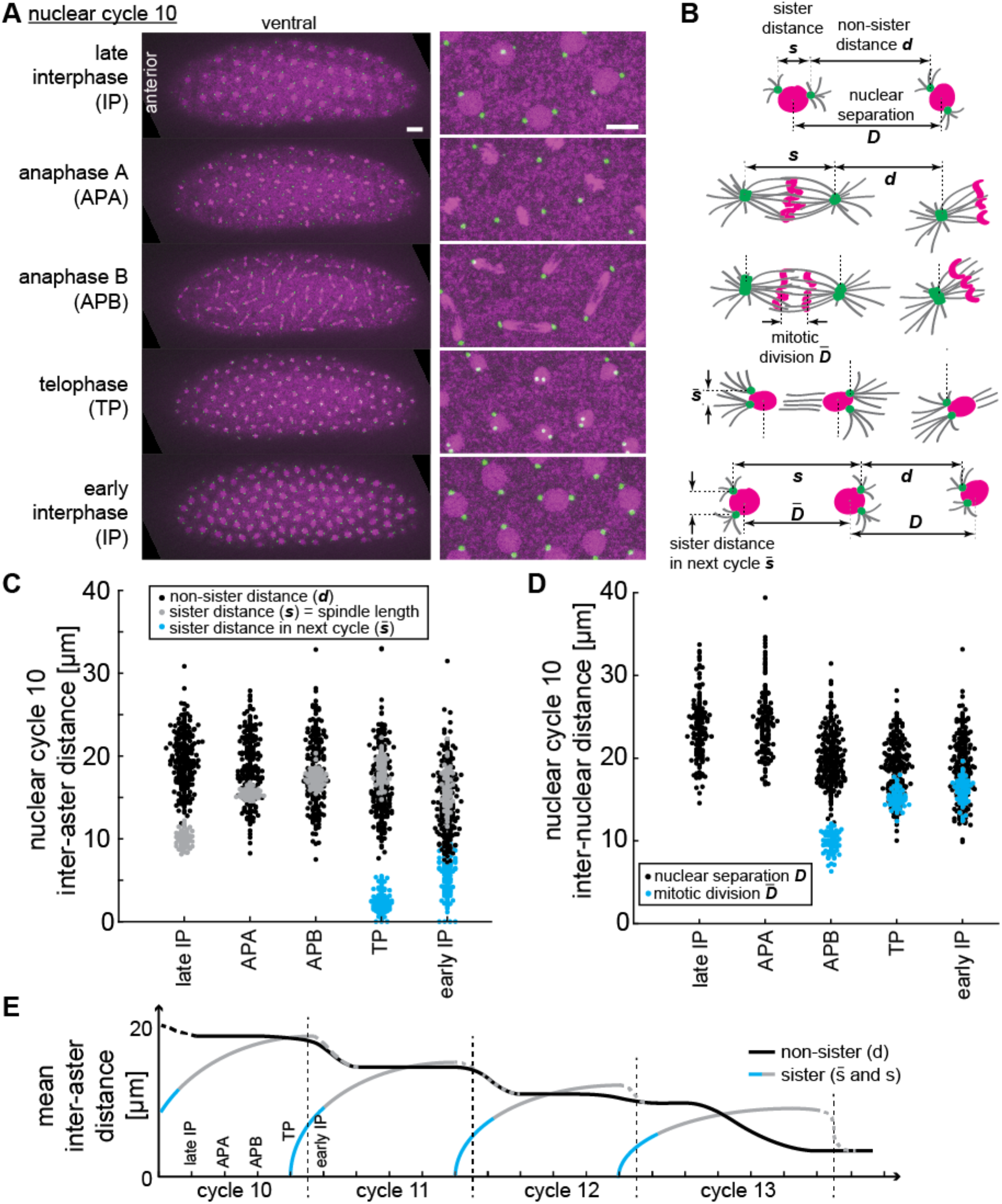
Spindle separation and orientation in the syncytial blastoderm of *Drosophila melanogaster* reveals robust distance maintenance. **(A)** Time lapse maximum intensity Z-projections and zoom-in images (right) from an embryo expressing H2Av::RFP (magenta) labelling chromatin and Spd2::GFP (green) marking centrosomes during nuclear cycle (n.c.) 10. Each panel corresponds to the indicated mitotic phase. Scale bars, 20 μm left, 10 μm right. **(B)** Morphological identification of mitotic phases and hierarchical classification of various distances between centrosomes or between nuclei. The scheme shows one nucleus assembling into a spindle, and one representative neighbour nucleus along the same cycle, with associated centrosome-nucleated microtubule asters. Nuclei in magenta, centrosomes in green, microtubules in grey. **(C)** Inter-aster distance during n.c. 10 (n=15 spindles, N=5 embryos). Neighbour distance *d* is shown in black, grey dots represent spindle expansion *s* and blue dots are sister centrosome separations 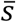 at spindle poles later in mitosis. **(D)** Corresponding inter-nuclear distance during n.c. 10. Blue dots show mitotic division of chromosomes. **(E)** Schematic of average inter-aster distances during blastoderm n.c. 10–13. Despite spindle elongation (grey lines) the neighbour inter-aster distance (black line) remains steady and decreases in early interphase of subsequent cycles. The extended dataset is presented in Suppl. Fig. 1.

There appears to be no long-range order in spindle orientation across the embryo^10^, but we asked whether there are local patterns. Defects may result in local ordering, akin to how subtle variations in sand corn size and shape induce fractures in sand piles with a range of lengths^32,33^. In the embryo, we investigated whether there were chains of aligned spindles (Fig. 2A–C and Methods). Calculating the probability of a given chain size (where size is defined by the number of nuclei belonging to the chain), *L*, in different cycles (Fig. 2D–E, Suppl. Fig. 2) we find *P*(*L*)~*L*^−*α*^, with *α* ≈ 2.0 *±* 0.3 in n.c. 13 (compared to *α* ≈ 3.6 *±* 0.5 with randomised spindle orientation). The value of *α* did not appear to decrease with increasing nuclear density (Fig. 2E), though it was dependent on the thresholds for defining chains (Methods). The value for *α* is similar to the exponent in cluster size variability in a range of other physical models^34–36^. Our results support the presence of short-range interactions driving spindle alignment in the absence of a membrane compartment.

**Figure 2 –.**
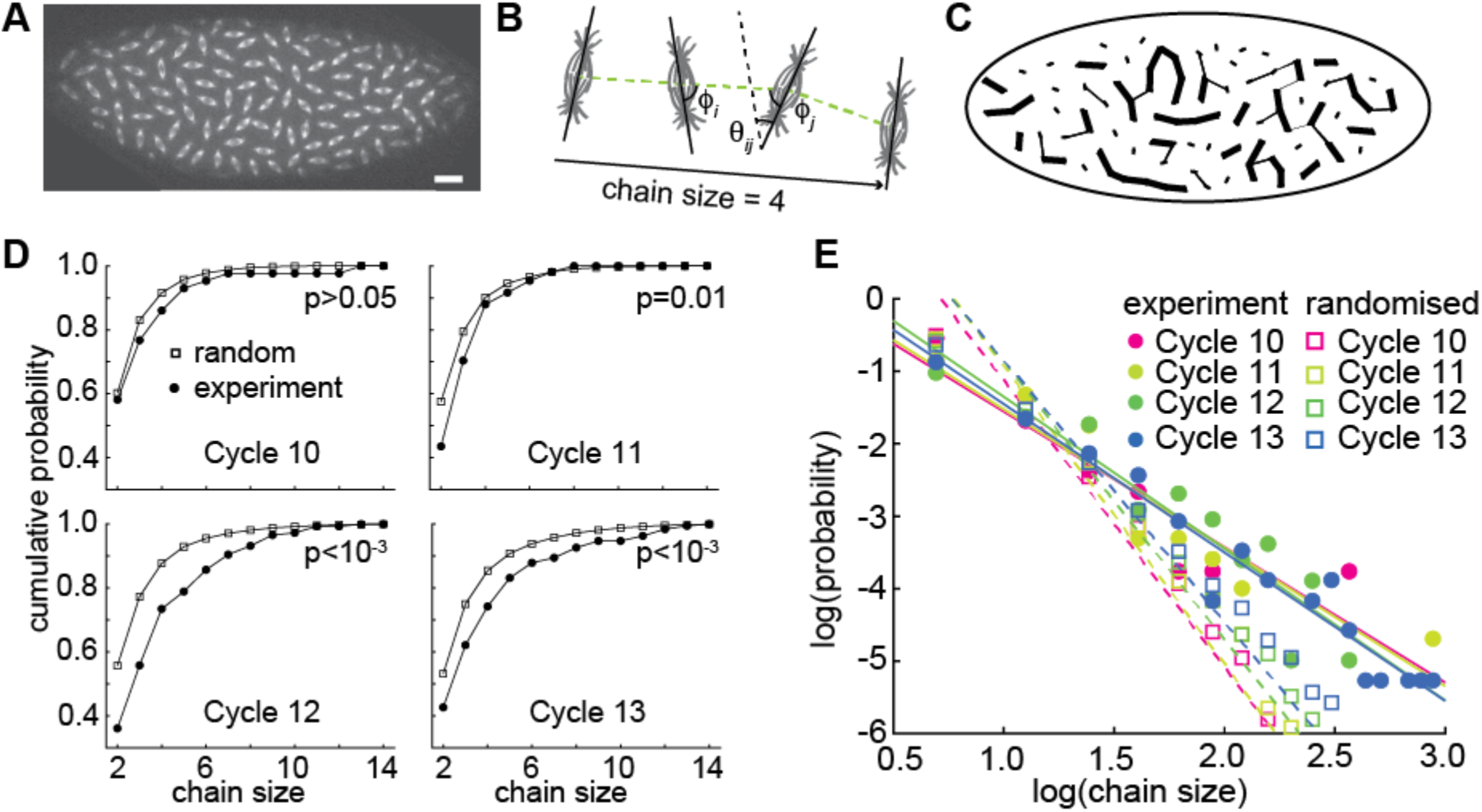
Local patterns of spindle orientation in the syncytial blastoderm indicate the existence of short-range interactions. **(A)** Maximum intensity Z-projection of an embryo in n.c. 11, expressing Jupiter::mCherry marking metaphase spindles (scale bar, 20 μm). **(B)** The schematic illustrates neighbouring spindles belonging to an alignment (‘force’) chain with size (number of members) *L* = 4. Spindles form angles *θ* and *ϕ* relative to each other (details in Suppl. Fig. 2A). Spindle alignment conditions were defined for *θ* (weak alignment) or for both angles (strong alignment). See Methods for details. **(C)** Resulting alignment chains for the image shown in **A**; the lines denote connections that satisfy two (thick) or only one (thin) of the conditions defining a chain. **(D)** Cumulative probability function of chain size for different cycles (n=7 embryos each for n.c. 10, 11, 12, 13). The p-value was calculated from Kolmogorov-Smirnov test. **(E)** Scaling of chain size probability with chain size for n.c. 10–13. Lines represent the fit to *βL^−α^*, where *L* is the chain size and a is the scaling exponent. Fitting parameters for all conditions are presented in the Methods. See also Suppl. Fig. 2.

We posited that repulsive interactions between astral microtubules underlie the forces determining the magnitude and spatial extent of the interactions defining spindle position and orientation. In this regard, the syncytium contrasts with uninuclear systems, in which spindle orientation is largely defined by cell geometry^29,37^. Direct interaction or fusion of astral spindles is inhibited by cell membrane boundaries formed during cytokinesis^38,39^. Here, we consider a simple model of aster-aster interaction in the shared cytosol of the syncytium (Fig. 3A): (i) asters have a radial microtubule structure nucleated from a centrosome; (ii) asters are self-repulsive due to microtubule interactions, generating a “dumbbell”-like potential for each nucleus; and (iii) nearest neighbour interactions dominate over longer-ranged interactions. Microtubule interactions can be mutual or mediated by molecular crosslinking^40^. Each spindle has a single rotational degree of freedom (Fig. 3A, bottom). From our model (Methods), we predict (Fig. 3B): (i) two nearby isolated spindles will align in parallel and orthogonal to their connecting line (i.e., *ϕ*~90°); (ii) three equidistant spindles align at *ϕ*~60° to each other; (iii) four and more spindles align randomly. The system is called “geometrically frustrated” for n>3 since multiple spindle configurations result in the same energy minimum^41^.

**Figure 3 –.**
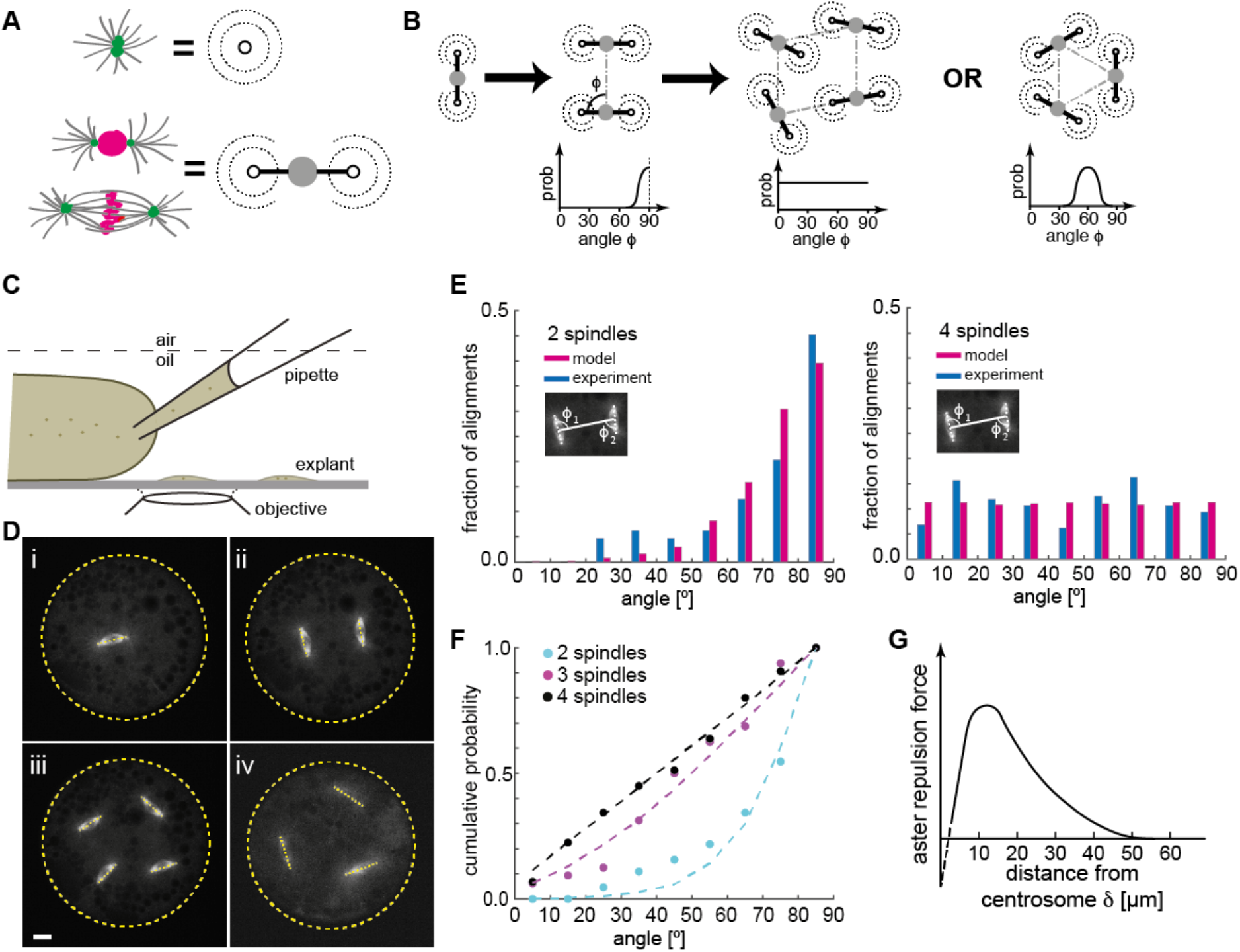
Division axis orientation for 2–4 spindles in a cytosolic explant are consistent with a simple dumbbell model of aster repulsion. **(A)** Schematic of an aster polymerised and organised by a centrosome (green), which translates into a concentric repulsion potential, here represented by dashed circles. When two asters are coupled to a nucleus (magenta) the repulsive potentials translate into a dumbbell potential with a rotational degree of freedom. **(B)** Scheme of dumbbells representing spindles in consecutive divisions. The graphs below show the expected probability of division angles *ϕ* between dumbbell axes from free energy considerations and assuming stochasticity (Methods). Right panel shows predicted alignment for a three-spindle arrangement. **(C)** Schematic of cytosol extraction from a *Drosophila* syncytial embryo and explant formation. **(D)** Maximum intensity Z-projections of explants from embryos expressing Jupiter::GFP (grey) and H2Av::RFP (not shown) containing different numbers of spindles. Dashed lines represent spindle axes, and yellow dashed circles represent explant boundaries. Scale bar, 10 μm. **(E)** Angle between the division axes of two spindles (left, n=32 explants) and between nearest neighbour spindles in a four-spindle scenario (right, n=40 explants) as measured in experiments (blue) or obtained from simulation (magenta, Methods). **(F)** Cumulative probability of the angle between division axes for two-, three- (n=5 explants) and four-spindle scenarios. Dashed lines are model predictions. **(G)** Proposed repulsion force as function of distance from the centrosome (green in **A**) with peak at ~15 μm and negligible for >45 μm. The dashed line represents the short-distance interaction regime that is below the diffraction limit of optical resolution. See also Suppl. Fig. 3.

In embryos, with hundreds of spindles, these predictions for small systems cannot be tested experimentally. Thus, we took advantage of our embryo explant assay, which enables us to study a small number of spindles in quasi-2D spaces^24^ (Fig. 3C, Suppl. Video 1). We measured spindle axis orientation *ϕ* relative to the separation axis (Fig. 2B). When a single spindle in an explant divides (Fig. 3D(i)), the two subsequent spindles align in parallel (Fig. 3D(ii)). Further divisions resulted in random spindle orientation, even when the spindles were uniformly distributed (Fig. 3D(iii), Suppl. Video 1). For three spindles, the alignment was biased away from random (p=0.04, Suppl. Fig. 3A–B), with some arrangements showing 60° alignment (Fig. 3D(iv), Suppl. Fig. 3C). However, we also observed cases where two out of three spindles aligned at 90°, likely due to the spindles not being positioned equidistantly. The experimental observations were in agreement with our model predictions when stochasticity was introduced for spindle orientation (Fig. 3E–F, Suppl. Fig. 3D–E and Methods). To conclude, in the absence of membrane boundaries the orientation and alignment of two-, three-, and four-syncytial mitotic spindles can be described by a model of mechanical dumbbells with nearest-neighbour repulsive interactions.

Centrosomes are essential for nuclear organisation in the embryo^21^. We posited that centrosomes, rather than nuclei, are the active positioning structures in the early embryo. In such a case, the aster repulsion is proportional to microtubule density and likely decays rapidly with distance *δ* from the MTOC at larger distances (Fig. 3G). To test these ideas, we utilised *giant nuclei* (*gnu*) mutant embryos, which undergo DNA endoreplication without mitosis; chromosome segregation is inhibited, leading to one or few polyploid nuclei, while centrosomes continue to duplicate and separate^25,26,42^ (Suppl. Video 2). We produced embryo explants from *gnu* mutant embryos and studied the positioning properties of a small number of microtubule asters in quasi-2D spaces. Asters consistently moved towards the centre of the explant, even when initially located near the boundary after cytosol deposition (Suppl. Video 3). We measured the radial intensity profile of single asters in explants as a proxy for aster size and microtubule length (Suppl. Fig. 4A–B). Away from the MTOC, the distribution was well approximated with a mono-exponential decay with decay length of ~12 μm (Suppl. Fig. 4B). This value is in excellent agreement with the size of asters associated to telophase and early interphase nuclei of wildtype embryo explants^20^. We conclude that, from the point of view of microtubule length regulation, *gnu* embryos mimic early interphase asters in wildtype embryos.

First, we explored the motion and positioning of single asters within our extracts. We measured the shortest distance of the centrosome from the boundary at steady state, which varied between *R*/2 and the maximum distance *R* (Fig. 4A). Deviation from precise centring may be due to yolk or lipid droplets (green circles in Suppl. Fig. 4A and Suppl. Video 3) forming exclusion zones. Typically, individual asters appear to self-centre within a restricted space, consistent with a radial force potential. We confirmed that our simple model of repulsive aster interactions (Methods) was able to replicate this observation (Suppl. Fig. 4C). Such short-ranged centring forces could be generated by microtubule polymerisation acting against the boundary of the water–oil interface^43,44^. Alternatively, hydrodynamic drag caused by microtubule motor transport together with radial asymmetry of asters can generate a net pulling force towards the centre^30,45^. Furthermore, there may be interactions between the aster and the yolk droplets present in the extract (see below). In these scenarios, once radial symmetry is restored the net force drops to zero.

**Figure 4 –.**
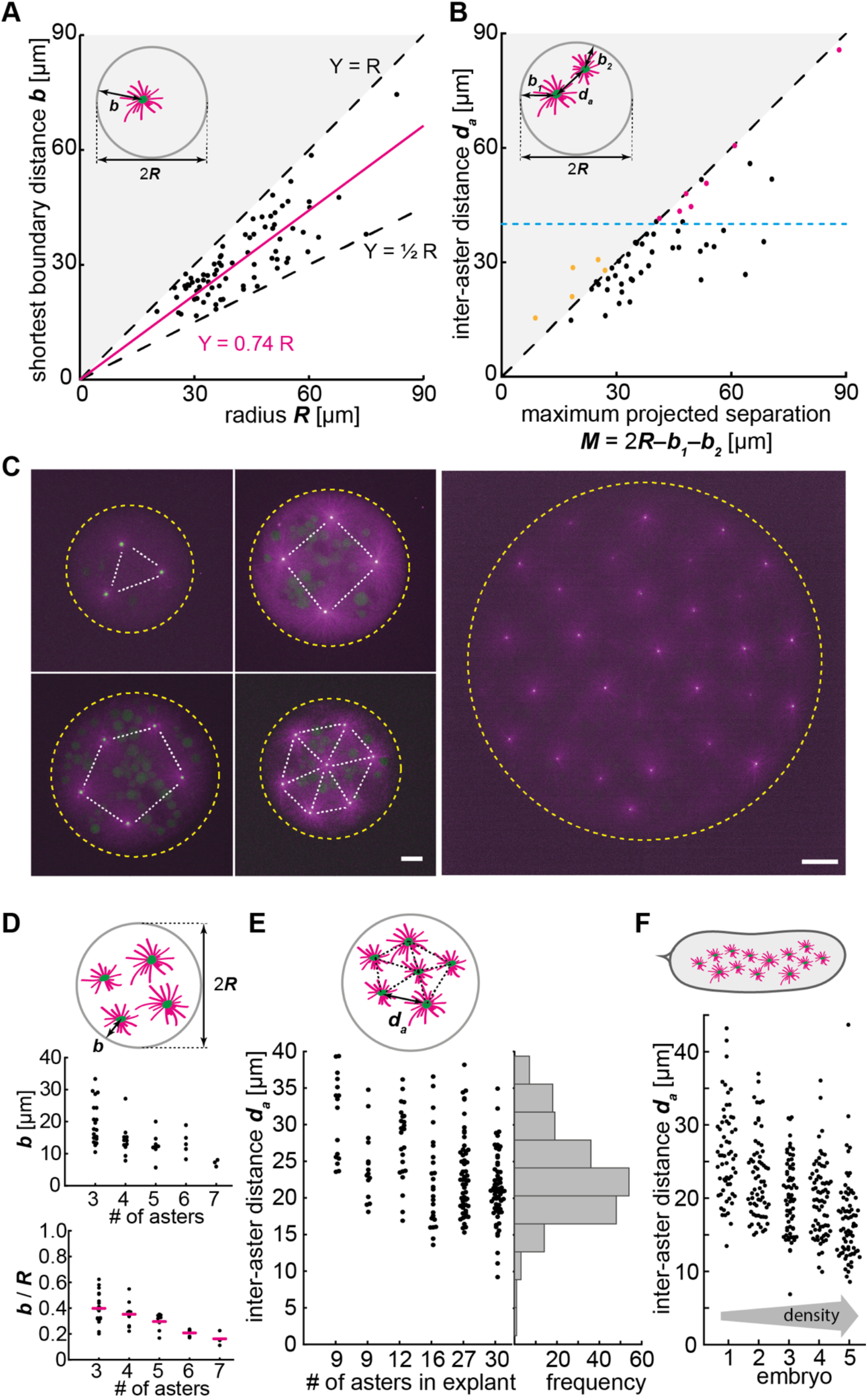
Free asters separate and achieve steady-state distance regularity. **(A)** Explant with single aster: scatter plot of the shortest distance to the boundary (*b*, see inset) as a function of the explant radius (*R*) (n=78). Magenta line: linear regression with zero intercept. **(B)** Explant with two asters: scatter plot of inter-aster distance (*d_a_*, see inset) as a function of the maximum projected separation *M*, calculated from the explant diameter (2R) and the boundary distance of each of the two asters (n=54). The blue dashed line represents the estimated upper limit of the interaction distance between two asters (~45 μm). Red dots represent cases where the two asters were positioned far apart during explant generation. The yellow dots are cases of small explants where projection leads to overestimation of *b* (Methods) **(C)** Maximum intensity Z-projections of explants containing multiple asters extracted from *gnu* mutant embryos expressing RFP::β-Tubulin (magenta) and Spd2::GFP (green) (scale bar, 10 μm). Dashed yellow circle represents the explant boundary, and white dashed lines highlight the symmetry in the aster distribution. **(D)** Distribution plot of shortest boundary distance (*b*, top scheme) and the ratio *b/R* from explants containing 3 (n=19), 4 (n=11), 5 (n=8), 6 (n=5) and 7 (n=3) asters. Magenta bars represent mean value. **(E)** Distribution plot of inter-aster distance (*d_a_*, see inset) of single explants containing nine or more asters. **(F)** Scatter plots of inter-aster distance from five *gnu* mutant embryos in order of increasing aster density. See also Suppl. Fig. 4.

How does aster positioning change in the presence of more than one aster? We investigated the steady-state distribution of two-aster configurations in explants. Inter-aster interaction, if existing, must balance with the force involved in moving asters away from the boundary. Two asters reached a steady-state separation distance that scaled with explant size and boundary distances, typically up to 45 μm, but did not scale further in larger explants (Fig. 4B). Consequently, the shortest boundary distances were not always diametral (Fig. 4B, inset). For most explants the inter-aster distance and the shortest boundary distances were similar (Suppl. Fig. 4D–F). Interestingly, two asters did not separate according to equal force but approximately partitioned the available space (*d* = *b*_1_ = *b*_2_ = 2R/3). Within our aster repulsion model, such a distribution could be replicated by having the repulsion from the explant boundary being larger than the aster-aster repulsion (Suppl. Fig. 4G, Methods). Some care is needed here though as (i) not all experiments covered the entire time course of aster separation; and (ii) some asters were likely initially positioned farther apart by the extraction procedure (Fig. 4B, red dots). Combined, these observations further support the presence of short-ranged repulsive interactions between asters and between aster and boundary. The two asters may mechanically interact via crosslinking of microtubules overlaps^40,46–48^, while astral microtubules may simply hit against the boundary interface, which acts as an immovable hard wall.

In explants, as the number of asters further increases, the shortest distance between the aster and boundary decreases (Fig. 4C–D). Interestingly, their steady-state position often assumed highly ordered, almost crystalline configurations (Fig. 4C), which we could recapitulate with our model (Suppl. Fig. 4H-I). Explants with higher numbers of asters had reduced aster separation distances, likely due to increased internal compression of larger 2D aster networks (Fig. 4E). Finally, in *gnu* mutant embryos, asters organise into a regularly spaced network and reach a dynamic equilibrium once the cortex becomes fully occupied (Suppl. Video 2). Quantifying inter-aster distance revealed a surprisingly stable and reproducible pattern along time (Suppl. Fig 4J–K). Notably, the inter-aster distance *d_a_* in *gnu* mutant embryos (Fig. 4F) at steady-state and in multi-aster explants are comparable, exhibiting density dependence, and matching the non-sibling distance *d* observed in wildtype embryos along division cycles, *i.e*. nuclear density (Fig. 1C,D). This indicates that positional patterning of centrosome-nucleated microtubule asters in the syncytial embryo may occur largely independently from embryo cortical factors, nuclei, spindle assembly and mitotic regulation.

Next, to probe the underlying mechanical interactions, we quantified the aster dynamics. Single asters located near the boundary after cytosol deposition stayed for up to 10 min, but they always eventually migrated (Fig. 5A, Suppl. Fig. 5A & Video 3, left). Single asters moved rapidly after separation from the boundary, with a maximum velocity of 0.05±0.02 μm/s, at around 20% of its final distance from the explant boundary (Fig. 5B, Suppl. Fig. 5B), before linearly decelerating and stopping between 15-35 μm from the boundary (Suppl. Fig. 5C). We noticed fewer microtubules oriented outwards when the aster was near the boundary (Fig. 5C), suggesting that most existing microtubules or those growing from the centrosome buckle and orient outwards, or they depolymerize rather than stabilise at short length. Indeed, in some samples we observed splay of microtubules near the explant boundary (Suppl. Fig. 5D). Single aster movement from the explant boundary could be reproduced with our dynamic model of aster repulsion, accounting for boundary effects (Suppl. Fig. 5E, Methods). There was only weak correlation between the final aster position and explant size, perhaps due to steric effects from lipid droplets (Suppl. Fig. 5A, inset). In summary, single asters display distinct dynamic phases, first as they separate from the edge and secondly as they migrate towards the explant centre.

**Figure 5 –.**
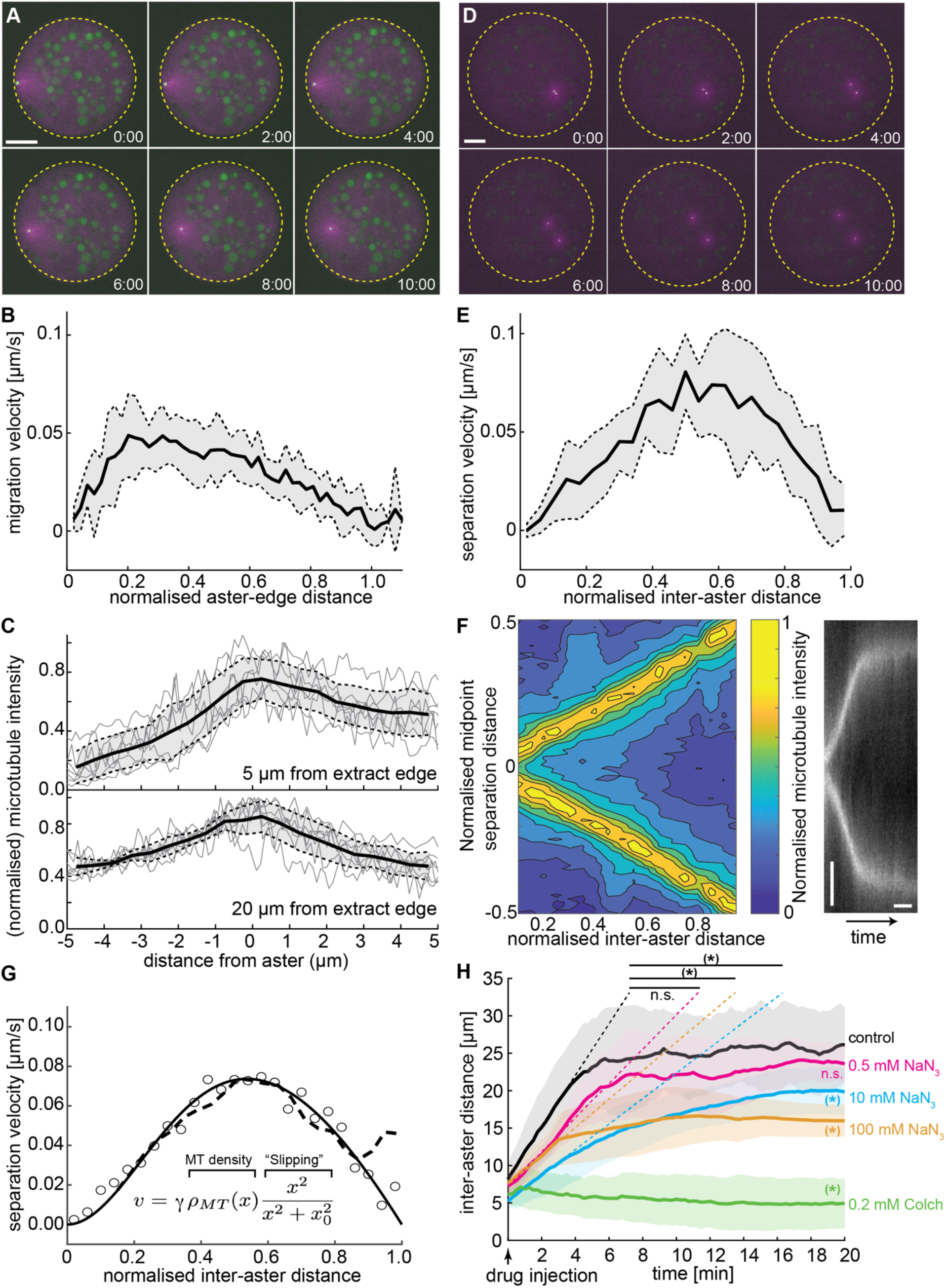
Aster dynamics in explants depends on microtubule distribution and interactions. **(A)** Maximum intensity Z-projections of a single aster moving away from the boundary of an explant produced from *gnu* mutant embryos expressing RFP::β-Tubulin (magenta) and Spd2::GFP (green). Yellow dashed circles represent the explant boundary. Scale bar, 20 μm. **(B)** Average migration velocity of single asters away from the explant boundary (n=7). Distance normalised by the final, steady-state distance for each aster (see also Suppl. Fig. 5A). **(C)** Average microtubule density (black line, inferred from RFP::β-Tubulin signal) along shortest distance to explant boundary, normalised by the maximum intensity within each experiment. Grey traces are individual experiments (n=7). **(D)** As in **A** but for an explant containing two separating asters. **(E)** Aster separation velocity as a function of normalised separation distance (n=9). For each experiment, distance is normalised by the final, steady-state separation distance (all data in Suppl. Fig. 6). **(F)** Left: Colourmap of normalised microtubule density between two separating asters (normalised as in **C**). Right: Kymograph of microtubule intensity between the asters during separation. Scale bars, 2 min (horizontal) 5 μm (vertical). **(G)** Fitting to average separation velocity (circles) considering microtubule intensity and a microtubule slipping term (inefficient repulsion). Microtubule density was either fitted beforehand (solid line, Suppl. Fig. 6E) or directly included (dashed line). **(H)** Aster separation dynamics upon injection of buffer (control, n=3), 0.5 mM (n=3), 10 mM (n=4), 100mM (n=3) sodium azide, or 0.2 mM (n=3) colchicine. * denotes p<0.05. Grey or coloured areas around average curves in **B**, **C**, **E** and **H** denote ± 1 s.d. See also Suppl. Figs. 5 and 6.

We further analysed the dynamics of the lipid droplets that are also present in the explant, to infer about passive behaviour from hydrodynamic effects. There was a droplet exclusion zone of ~10 μm around each aster. As the aster moved away from the boundary, the lipid droplets streamed around the aster, maintaining their exclusion (Suppl. Fig. 5F–G). We quantified the motion of lipid droplets with and without an aster present (Suppl. Fig. 5H). In the absence of an aster, the lipid droplets appeared to move randomly (*x_rms_ ~t*^1/2^). In the presence of an aster, lipid droplets moved faster, and appeared to move in a more directed manner (*x_rms_ ~t*^2/3^). Such behaviour is consistent with the aster having a repulsive force potential that can act on the surrounding lipid droplets and other boundary constraints.

To gain information about the aster-aster interactions, we next tracked aster pairs during separation (Fig. 5D–E, Suppl. Fig. 6A–D & Video 3, right). The final distance between asters correlated well with explant size within sizes tested (Suppl. Fig. 6A). Interestingly, the peak separation velocity was always near half the final aster separation distance (Fig. 5E), independent of final separation distance. This contrasts with the single aster scenario (Fig. 5B), suggesting that the overall effective forces are different in the two cases, consistent with our observations in Fig. 4D–E. Given this eccentric movement, the aster separation could be driven by overlap and sliding of astral microtubules^17,48,49^, or by mutual contact leading to repulsion by microtubules of both asters. Thus, we quantified the microtubule intensity between the separating asters (Fig. 5F, left) and generated kymographs of the microtubule fluorescence intensity along the separation axis (Fig. 5F, right). The intensity at half the separation distance decayed exponentially (Suppl. Fig. 6E), consistent with models of dynamic microtubule length distribution^50,51^. When aster separation ceased there was almost no measurable microtubule signal between the asters.

For a viscous material, the velocity, *v*, of an object is dependent on the applied force *F: v ≈ γF*, where *γ* is the effective viscous drag coefficient. Naïvely interpreting the microtubule distribution as the resulting force profile does not match with the observed separation velocity profile. However, multiplying the microtubule distribution by an effective *slipping* term, 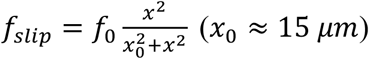, results in an excellent fit to our observed aster separation velocities (Fig. 5G). We attribute this effective slipping to molecular friction between microtubules^52^. Implementing such a force profile within our model of repulsive asters, we were able to qualitatively replicate the observed experimental observations (Suppl. Fig. 6F). Finally, we noticed that the microtubule density between the asters was often not maximal along the shortest distance between the asters (Suppl. Fig. 6G–H), suggesting that the contact interfaces between asters is more complex than assumed above. Overall, we see that aster-aster and aster-boundary dynamics both appear to involve repulsive interactions with effective slipping at very short distances, though the aster-aster interactions are weaker than those between the asters and the boundary.

To further explore the nature of the aster interaction, we performed a series of inhibitory treatments. Since small-molecule inhibitors for candidate molecular motors have no effect in *Drosophila*^53,54^, we targeted microtubules and ATPases in general. We generated explants with two asters in the course of separation and pulse-injected a defined volume of 200 μM colchicine, which causes acute depolymerisation of microtubules. Upon injection the asters stopped separating and sometimes inverted their direction of motion (Fig. 5H, Suppl. Video 4). We then tested whether active molecular machinery was required for aster repulsion by inhibiting ATP consumption. We injected a series of concentrations of sodium azide into explants that contained a separating pair of asters (Suppl. Video 4). Adding sodium azide decreased the initial recoil velocity (dashed lines in Fig. 5H) and also resulted in a considerable reduction in aster separation. However, even at very high concentrations of sodium azide, we still observed residual motion, suggesting that both actively driven microtubule-mediated separation and passive physical contact driven separation occur.

From our observations, we conclude that aster separation is caused by a net repulsive force between asters and between aster and boundary. However, either overall pushing or pulling can cause the separation dynamics and centring^31,55,56^. In particular, pulling within the cytoplasm requires aster asymmetry^30,45^. Thus, we performed targeted UV photo-ablation experiments in explants and generated ellipse-shaped ablations positioned asymmetrically around one aster, affecting microtubules on the left side more than on the right side of the aster (Fig. 6A, Suppl. Video 5). If pulling on the boundary^31^ or hydrodynamic effects from vesicle transport along microtubules^30^ drives aster motion, we expect a displacement to the right (positive) after ablation. Conversely, if the net force applied on microtubules favours pushing on MTOC, we expect a displacement to the left (negative). Indeed, asters consistently moved to the left, supporting a dominating effect of microtubule-driven pushing (Fig. 6B). As a control, we performed the same perturbation in explants that were injected with the microtubule inhibitor colchicine (Fig. 6C). Under this condition, asters moved very slowly to the right (positive), which is consistent with a weak hydrodynamic effect from other contractile sources (*e.g*. actomyosin^57^). We conclude that a single aster moves and positions within explants by microtubule-dependent pushing force.

**Figure 6 –.**
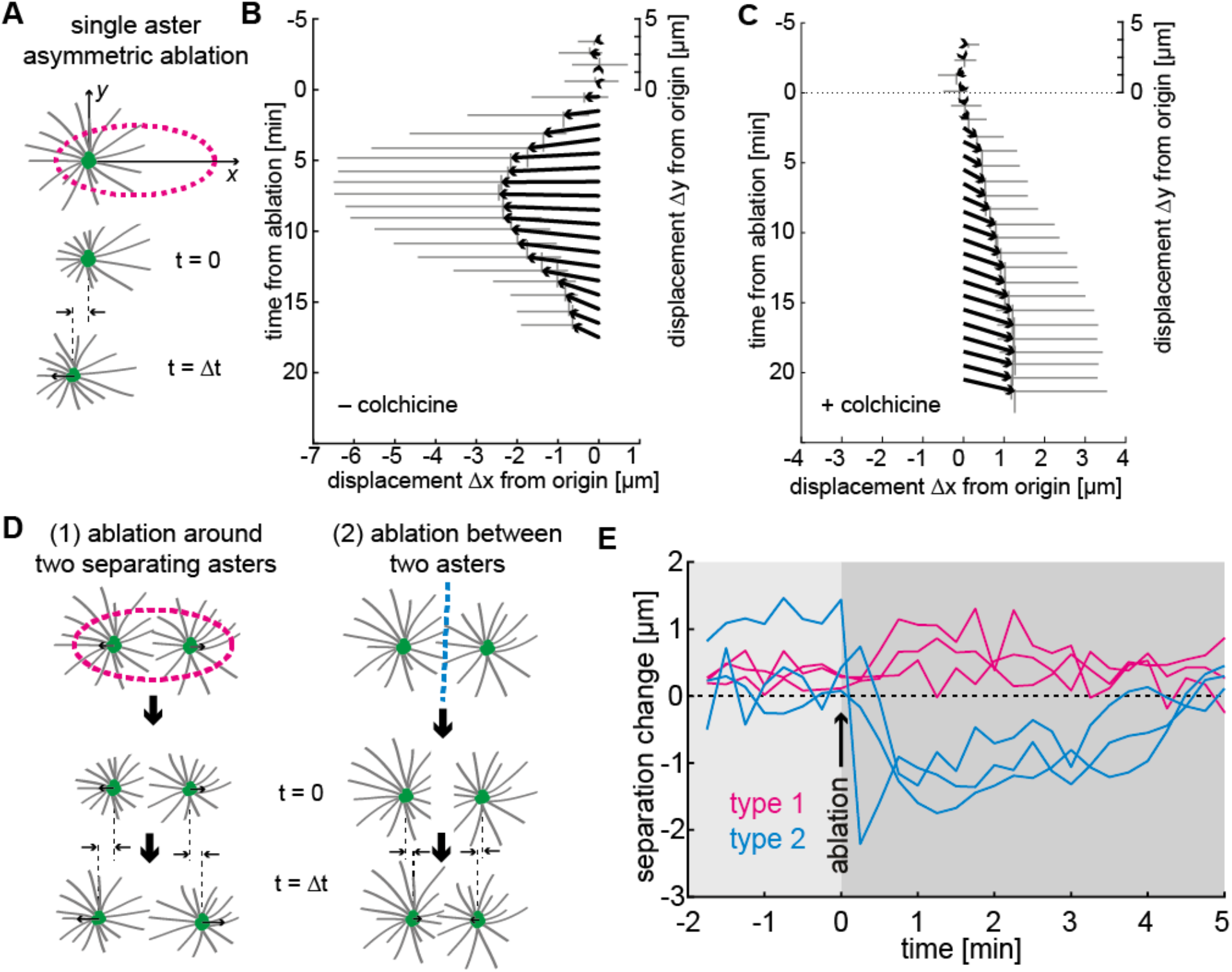
Aster positioning and separation is determined by a dominant microtubule-dependent pushing force. **(A)** Schematic of single aster eccentric circular UV laser ablation (magenta dashed line); this ablation aims at shortening astral microtubules on the left side of the aster. t=0 min denotes ablation time. **(B–C)** Aster displacement before and after eccentric circular ablation in explants unperturbed (**B**, n=8) or treated with colchicine (**C**, n=8). Arrows represent average displacement magnitude and direction, and vertical and horizontal grey bars denote ±1 s.d. of displacement in x and y, respectively. **(D)** Explants containing two asters were perturbed by (1) ellipse ablation around both asters during separation (“peripheral ablation”); (2) linear ablation between two asters (“central ablation”). **(E)** Change of inter-aster distance upon laser ablation (time = 0) as described in **D**. Upon peripheral ablation, separating asters maintained their movement and sometimes slightly accelerated, while central ablation caused movement towards each other. See also Suppl. Video 6.

To challenge these conclusions, we performed two types of ablation in explants containing two asters (Fig. 6D, Suppl. Video 6): 1) light pulses emitted along an ellipse around both asters to destroy microtubules in the periphery; 2) light pulses emitted along a line between the two asters to destroy microtubules between asters. If forces are attractive, then ablation type 1 will stop separation while ablation type 2 will lead to an acceleration. If forces are repulsive, we predict the opposite response. We found a slight acceleration for peripheral ablation and a strong deceleration with recovery for central ablation (Fig. 6E). Separation recovered likely because of fast regrowth of microtubules after ablation (in the range of μm/min^58^). In summary, the dynamic behaviour of asters in our explants is consistent with a model of radially symmetric microtubule-based repulsion.

What is the relevance of our findings *in vivo?* Specifically, do these aster interactions enable the embryo to pack the nuclei in a regular manner? Heterogeneities in nuclear density are a common phenomenon in early embryos resulting from aberrant cortical migration or nuclear internalisation due to mitotic failure (Fig. 7A, Suppl. Fig. 7A & Video 7). We predicted that nuclei neighbouring a low-density region will orientate their division axis towards that region, where repulsion is weakest. We identified such low-density regions and quantified the subsequent division orientation of the surrounding nuclei, which confirmed our prediction (Fig. 7B, Suppl. Fig. 7B). Repeating the same analysis on regions of uniform nuclear density showed no correlation in the division angle (Suppl. Fig. 7C). We also generated acute density reductions by UV ablation. Using low laser damage, the target nuclei failed to divide and detached from the cortex lowering local nuclear density (Suppl. Fig. 7D–E). Subsequently, the surrounding nuclei adjusted their division axis to orientate into the perturbed region (Suppl. Fig. 7F). Combining our results from spontaneous low-density regions and laser-ablated embryos (including larger ablations of 3–5 nuclei), we see that the microtubule repulsion mechanism is efficient in adjusting the angle of division to compensate for heterogeneities in nuclear packing (Fig. 7C).

**Figure 7 –.**
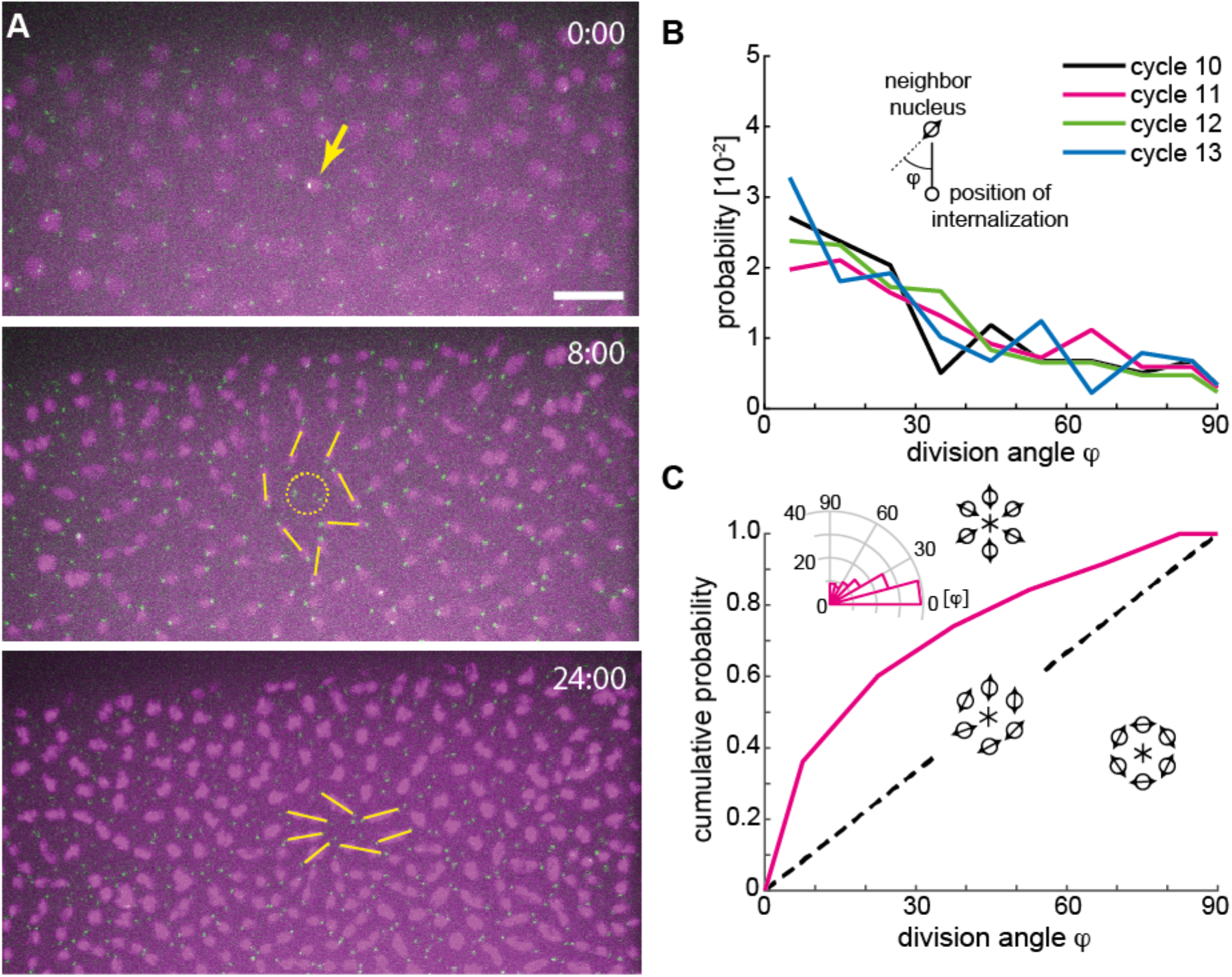
Microtubule-dependent repulsion provides a mechanism for spindle alignment towards lower density. **(A)** Maximum intensity Z-projections from an embryo expressing H2Av::mCherry (magenta) and Spd2::GFP (green) in n.c. 12–13. Yellow arrow (top panel) denotes internalisation of a nucleus. The centrosomes remain at the embryo cortex (yellow circle, middle panel). Division axes of neighbouring spindles (yellow lines) orientate towards the location of internalisation in n.c. 13 (bottom panel). Scale bar, 20 μm; time in min:sec. **(B)** Probability density function of the division angle orientation *φ* of neighbouring nuclei to regions of low nuclear density in n.c. 10–13 (n=67, 116, 96, 73 angles from N=10, 15, 15, 15 embryos in n.c. 10, 11, 12, 13 respectively). **(C)** Cumulative distribution function of division axis angle at the end of n.c. 13 towards artificially generated holes generated by single-pulse UV laser ablation (n=108 angles from 15 embryos). The dashed black line represents random division orientation. See also Suppl. Video 7, and extended data in Suppl. Fig. 7.

## Discussion

Robust embryonic development critically depends on homogenous delivery of nuclei to the cell cortex and subsequent maintenance of a regular nuclear distribution despite further division cycles^15,16^. Recent work has shown how nuclear divisions are synchronised and, as a consequence, how nuclei are globally distributed around the embryo cortex^4,12^. However, these results assumed that nuclei are positioned regularly after each round of duplication. Here, we asked whether we could understand the mechanical circumstances ensuring such local order of nuclei. To answer this requires deepening our understanding of the biophysical principles defining the orientation of the spindle axis and how nuclei separate and reposition during division cycles. Our explant experiments demonstrate that, in the absence of perturbation by neighbour interactions, the ground-state orientation of a spindle is orthogonal to the previous division axis (Fig. 3F). This did not depend on the size of the explant and we found the same pattern in very large (>200 μm) or small explants. We conclude that geometry has little or no effect on the orthogonal sequence of division axes in pseudo-2D spaces^37^. Our results can be explained by the stereotypical migration of the two centrosomes from their common origin, each along one quadrant of the nucleus, until they form the poles of the bipolar spindle^59^. Hence, orthogonality of spindle axes likely emerges from the geometric nature of bipolar structures and symmetry considerations. Surprisingly though, a system of four or more spindles in a two-dimensional space evolves towards random orientations, arguing against active spindle orientation control by the cell. Our analysis suggests that force balancing and energetic minimisation in a noisy two-dimensional environment dynamically determine where nuclei are positioned and in which orientation they propagate upon division. Interestingly, in larger networks, small positional irregularities result in division axis orientation towards the low-density region, enabling the nuclear distribution to homogenise quickly and act as a self-repair mechanism. Finally, it would be interesting to compare aster force driving spindle alignment in the early embryo with similar microtubule-driven processes across cells, such as mitosis in polarised tissue growth^60^.

Dissecting the molecular mechanism of microtubule aster repulsion in the embryos by genetic manipulation is challenging since many microtubule-associated proteins and motors play an essential role during oogenesis and early embryogenesis. Moreover, unlike in other species, available small-molecule inhibitors do not specifically target these motors in *Drosophila*^53,54^. Alternative approaches have been used, such as antibody-mediated inhibition, TEV-mediated protein cleavage, or germline specific RNAi. Inhibition of Klp61F, a promising candidate for driving microtubule-based repulsion^17^, causes a strong spindle assembly phenotype in syncytial embryos^61^. In combination with knock-down of the antagonising Ncd (kinesin-14) spindle assembly was rescued but spindles and daughter nuclei failed to separate properly^61^. In a transgenic Klp61F null construct expressing TEV-Klp61F-GFP, nuclei were more disordered in interphase following injection of TEV, which chemically ablates the motor^48^. However, the authors doubted microtubule sliding of Klp61F being essential for nuclear positioning as they recorded higher mobility of nuclei after TEV injection. Recently, we have shown that Fascetto (Feo), a microtubule crosslinker of the PRC1/Ase1 family, and Klp3A (kinesin-4) colocalise as puncta in regions between neighbouring nuclei. Depletion of Feo leads to irregular delivery of nuclei to the cortex and loss of separation after nuclear repositioning by micro-manipulation^49^. Is the observed aberrant nuclear movement and positioning in the syncytial embryo of the above mutants due to pushing or pulling forces, and what role does the nucleus play? In the present study, we provide evidence that there exists mechanical repulsion between asters independent of the nucleus and cell cortex.

Aster positioning and spindle axis determination have been studied by cell and developmental biologists for several decades^62–67^ and have seen renewed interest in recent years due to the advent of new techniques in imaging, sample control and perturbation methods^28,29,68^. Aster positioning is important in egg and early embryo cells; it is at the core of pronuclear apposition after fertilisation and determines the cell division plane during early mitotic blastomere divisions^27,68–70^. In eggs and embryo cells from *C. elegans* or sea urchin, mitotic spindle positioning is likely controlled by cortical pulling forces^29,31^. However, this model does not explain observations in large cells in which repositioning occurs before astral microtubules contact the distal cell wall^27,38,71^. Sperm aster movement was initially believed to depend on cell wall pushing^69^. However, the mechanism appears to depend on cytoplasmic pulling, at the core of which is vesicle movement from the periphery towards the aster centre driven by cytoplasmic dynein^28,30,45^. In this scenario, the net force on the aster is dependent on astral microtubule length and, thus, on spatial asymmetry of microtubule density. Yet, this model is currently contested by experiments that maintain support of the pushing model^72^. Further, recent observations in *Xenopus* egg extract, either in combination with reconstituted cortical actin^73^ or exposed to artificial geometric constraints^74^, suggest that mechanisms exist for aster positioning beyond hydrodynamic pulling^75^. Cells appear to utilise a combination of possible mechanisms, depending on spatial circumstances and the process to be achieved, which then leads to a net pulling force or a net pushing force on the aster. Nevertheless, all these model systems have in common that cytokinesis ensures the cytosolic isolation of spindles, and neighbour interactions never occur. The mechanics of aster positioning in multinucleated cells is yet more complex, with a large array of possible interactions. This may be why aster mechanics have not been addressed in the *Drosophila* syncytium, otherwise a popular model system to study development. Here, we provide definitive evidence, using a reductionist approach, that a mechanism generating net repulsion between asters and towards the physical boundary has emerged, which robustly and homogenously distributes syncytial nuclei.

Why is a high spatial regularity of nuclei important for the embryo? After n.c. 13, the embryo transforms into a multicellular embryo by engulfing each nucleus with plasma membrane^1^. During this process, the nearest neighbour internuclear distance defines cell size. Therefore, a narrow distance distribution leads to a uniform size of cells that subsequently assume distinct function during body part definition. Analysis of information decoding in the *Drosophila* embryo has shown how each individual cell unambiguously reads its current position, which defines a specific function later in development^5^. But these results are dependent on the interpreting units (*i.e*. the nuclei) being uniformly distributed around the embryo. Therefore, we can conclude that a robust mechanism defining cell size and position is crucial as size irregularity would effectively decrease positional precision.

## Materials and Methods

### Fly strains

Flies with genotypes w^1118^; +; endo>Jupiter::GFP (stock no. 6836, Bloomington) and w*; +; endo>H2Av::RFP (stock no. 23650; Bloomington) were crossed to generate recombinant progeny. Similarly, flies expressing fluorescent reporters recombined on the 2^nd^ chromosome were produced by crossing the following stocks: w*; endo>H2Av::RFP; + (stock no. 23651; Bloomington); w*; pUbq> β-Tubulin::EGFP; + (stock no. 109603, Kyoto); w*; pUbq>RFP::β2-Tubulin; + (originally described elsewhere^76^) and w^1118^; pUbq>Spd2::GFP; + (all provided by Mónica Bettencourt Dias, Instituto Gulbenkian de Ciência, Portugal). All resulting recombinant fly lines are homozygous viable. w^1118^; +; Jupiter: :mCherry flies were generated by and obtained from Nick Lowe in Daniel St Johnston’s lab (The Gurdon Institute, United Kingdom). H2Av::mCherry flies were generated as described elsewhere^14^. Two different mutants of giant nucleus (*gnu*), namely w*; +; gnu^305^/TM3 (discontinued stock no. 3321; Bloomington) and w*; +; gnu^Z3-3770A^/TM3 (discontinued stock no. 38440; Bloomington), were each balanced with w^1118^; CyO/Sco; MKRS/TM6B (stock no. 3703, Bloomington). Above-described recombined lines on the 2^nd^ chromosome were individually crossed with gnu mutants and kept as balanced stocks. Finally, trans-heterozygous were generated for gnu^305^/gnu^Z3-3770A^ mutants, whereby only flies homozygous for the fluorescent reporters on the 2^nd^ chromosome were selected for increased signal collection during live microscopy. These trans-heterozygotes laid fertilized eggs which undergo several embryonic rounds of chromatin replication and centrosome duplication, allowing for the study and quantification of asters at the embryo cortex.

### Embryo collection and sample preparation

We followed established procedures^77^ of fly husbandry, keeping flies at 25°C under 50-60% humidity. For embryo collections, young adult flies were transferred to a cage coupled to an apple juice agar plate. After 2–3 rounds of egg laying synchronisation, developing embryos were collected every 30–60 minutes. In the case of *gnu* mutants, embryos were collected at different time intervals, ranging from 30 min up to 4h. Embryos were dechorionated by short immersion in 7% sodium hypochlorite solution (VWR). After extensive rinsing with water, embryos were aligned and immobilised in a thin strip of heptane glue placed on 22×22mm coverslips, and covered with halocarbon oil (Voltalef 10S, Arkema).

### Microscopy

Time-lapse acquisitions were conducted on a Nikon Eclipse Ti-E microscope equipped with a Yokogawa CSU-W Spinning Disk confocal scanner and a piezoelectric stage (737.2SL, Physik Instrumente). For embryo imaging, 15 μm (31 planes) Z-series stacks were acquired every 15s (wildtype, if not states else) or 30s (*gnu* mutant), using a Plan Fluor 40x 1.3NA oil immersion objective, the 488nm and 561nm laser lines, and an Andor Zyla 4.2 sCMOS camera to acquire images. For explants up to 100μm in diameter, we used a Plan Apo VC 60x 1.2NA water immersion objective with 2x post-magnification and an Andor iXon3 888 EMCCD camera. When needed, the Andor Zyla 4.2 sCMOS camera was selected to acquire a 2x wider field of view with the same spatial resolution or, alternatively, the Apo λ S LWD 40x 1.15NA water immersion objective. For acquisition in explants, the frame rate was 15s for *gnu* mutant 30 s for wildtype embryo explants.

### Single embryo explant assay

Embryo extractions were performed as previously described^24,78^. Briefly, cytosol from wild-type embryos between telophase and subsequent interphase of cycle 8 was extracted by puncturing the vitelline membrane with a sharp glass micropipette and flow control by operating a bi-directional syringe pump. Small explants of cytosol (in the picolitre range) were deposited on poly-L-lysine coated glass surface under halocarbon oil. Time-lapse acquisitions typically started in late interphase or prophase. In the case of *gnu* mutant embryos, most extractions were performed when few centrosomes (between 5 and 40) were visible at the anterior-lateral cortex. During extractions, shear stress was avoided to prevent structural damages and undesirable molecular dissociations that induce premature mitotic failures or aberrant microtubule structures. In *gnu* mutant embryos, repeated use of the same extraction micropipette is not recommended. Explants from wildtype embryos initially containing a single nucleus were selected for time-lapse imaging of subsequent mitotic divisions. Explants from *gnu* mutants initially containing a single free aster near oil interface or two free asters in close proximity were selected for time-lapse imaging of aster separation. All experiments were conducted at 25±1 °C.

### Pharmacological perturbation of embryo explants

Pharmacological perturbations were performed by adding different drugs (colchicine at 0.2 mM, sodium azide at 0.5, 10, or 100 mM) diluted in cytoplasm-compatible buffer (50 mM HEPES, pH 7.8, 100 mM KCl, 1 mM MgCl2). Solutions were directly administrated to the explants using a fine pipette (pulled using a Narishige PC-100 Puller with a: 2-step (69% + 55%) heating protocol and with 4 mm drop length) connected to an Eppendorf FemtoJet^®^ 4i pump. The final drug dilution in the explants was of approximately 1:10 (solution:cytosol). Buffer injections were conducted as control.

### Laser ablation system

The laser ablation systems used for experiments with intact embryos (at EMBL Heidelberg, described elsewhere^20^) and embryo extracts (at Instituto Gulbenkian de Ciência, implemented by IA Telley on the microscope described above) were conceptually identical. A Crylas FTSS-355-Q pulsed laser emitting 355 nm, 1.1 ns pulses, 15μJ pulse energy at 1 KHz was aligned with a beam expander (16x), a scan head (SCANcube 7, Scanlab, Germany) coupled to an f-theta lens (f=56 mm, anti-reflection coating for 340–370 nm, SCANLAB AG, Germany). The focus point of the f-theta lens was aligned to be parfocal to the focal plane of the objective, using a tube lens (f=200 mm, Ø=30 mm, 355 nm AR coated, OWIS, Germany) and a dichroic mirror (T387 DCLP, Chroma) in the upper stage filter wheel. Any scattered light was blocked at the emission side with a RazorEdge LP 355 dichroic mirror OD6 @ 355nm (Chroma). The system was controlled with homemade journals for Metamorph software (Molecular Devices Inc.). The optimal laser power was set to ensure microtubule ablation while avoiding thermal expansion of cytoplasm, with post-ablation microtubule signal recovery matching known polymerisation dynamics. This combination of conditions proved to be efficient at ablating target structures beyond fluorophore bleaching. In explants containing a single aster, astral microtubules were asymmetrically ablated by positioning an ellipsoid off-centre (21.7 by 10.8 μm, 4 times, 15s interval, 0.54 μm step, laser power: 25%) (Fig. 5A). In explants containing two asters, astral microtubules were ablated using an ellipsoid (21.7 by 10.8 μm, 3 times, 15s interval, 0.54 μm step, laser power: 10–15%) roughly centred at the mid-point between the two asters, while interpolar microtubules were ablated using linear ablations (21.7 μm, 3 times, 15s interval 0.54 μm step, laser power: 10–15%) perpendicular to the axis connecting the asters (Fig. 5D).

### Distance analysis in embryos

Automated positional detection of the signals from centrosomes and nuclei (or chromatin) was performed by applying a Gaussian blur filter (radius: 1–2 pixels) and using the plugin TrackMate v3.5.1 in Fiji ImageJ^79,80^. The coordinates of detected spots were imported into MATLAB^®^ for assignment and distance calculation. The connection between poles belonging to a spindle structure was assigned in a custom-made script requiring user input, on an area containing 15–40 spindles for each mitotic phase and per embryo. For each spindle–assigned coordinate positions, the nearest neighbour positions were determined using the Delaunay triangulation functions in Matlab^®^ yielding a connectivity list. Thereby, a spindle structure is defined as a combination of *n* centrosome and *m* chromatin positions (*n,m*) with the following numbers for mitotic phases: late interphase (2,1); anaphase A (2,1); anaphase B (2,2); telophase (4,2), early interphase (4,2). With this assignment, the duplicated organelles dissociate at the transition from telophase to early interphase, so that two related nuclei become independent neighbours. Next, the 3D Euclidean distances between relevant positions were calculated from position coordinates with a computer-assisted manual heritage classification. The distance between separating chromosomes 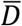 was calculated from the two chromatin entities within a spindle. Spindle length s was calculated from the distance between two centrosomes belonging to each spindle. In phases with four centrosomes per spindle, two at each pole, spindle length was defined as the smallest distance between opposite centrosomes (four possible combinations). The sister centrosome distance 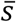 was calculated from centrosome pairs at each spindle pole. Inter-aster distance *d* (corresponding to the distance between non-sister centrosomes) was calculated between different, neighbouring spindles by selecting all centrosomes not associated with the same spindle from the nearest neighbour connectivity list. Inter-nuclear distance *D* was calculated between nearest neighbour nuclei or chromosomes not belonging to the same spindle. Finally, the arithmetic mean and standard deviation of the distance distributions within a single embryo were calculated and overlaid for division cycles 10–13 (Suppl. Fig. 1). In *gnu* mutant embryos, inter-aster distance *d_a_* was calculated from a selected region containing 15–20 centrosomes, for each embryo at different time points using triangulation and neighbourhood connectivity list. Centrosomes located at the anterior hemisphere were excluded to avoid the influence of the giant polyploid nucleus. The *gnu* mutants present variable centrosome densities depending on age and other unknown factors. We analysed the variation of distance distribution for five different embryos with similar densities during intervals of 10 min. All data plots were generated in MATLAB^®^.

### Spindle alignment analysis in embryos

For each embryo, we took the time point two minutes prior to the first onset of nuclear division. The angle of the centrosome pair was used to define an orientation angle for the spindle. In calculating the force chains, we did the following for each cycle:

1. For each spindle *i*, identify nearby spindles (taken as 1.3 times {average nearest neighbour distance});
2. Calculate the orientation angle difference between pairs of nearest spindles. This value can range between 0° and 90° (as there is no direction to the spindle orientation). We define this angle as *θ_i_* for each pair of spindles denoted by *i* and *j* respectively.
3. Find the vector between each spindle and its nearest neighbours **x**_ij_. Find the angle, ϕ, between the orientation of the spindles *i* and *j* relative to **x**_*ij*_. Notably, as the orientation does not have a specific direction, for angles >90° we define Φ = 180 − ϕ.
4. We tested different thresholds on θ and ϕ (annotated by Δ*θ* and Δ*ϕ*) to find the chain length frequency (Suppl. Fig. 2C). For a spindle to belong to a chain, at least the angle *θ* had to meet the condition 0 ≤ *θ* ≤ Δ*θ*. In addition, a second condition was defined: Δ*ϕ* ≤ *ϕ* ≤ 90°. Neighbouring spindles that meet both conditions belong to a chain that exhibits alignment both along the chain axis and between spindles. In the below table, we give the fitted value of a (as defined in main text) for different constraints on *θ* and *ϕ*.

**Table 1A:** α for different conditions on chain alignments for experimental data

| Cycle | $\theta = \phi = 45^\circ$ | $\theta = 45^\circ, \phi = 30^\circ$ | $\theta = 30^\circ, \phi = 45^\circ$ | $\theta = \phi = 30^\circ$ |
| --- | --- | --- | --- | --- |
| 10 | 1.9 | 2.7 | 2.7 | 3.4 |
| 11 | 1.9 | 3.3 | 2.9 | 4.0 |
| 12 | 2.1 | 3.1 | 3.4 | 3.8 |
| 13 | 2.0 | 3.4 | 3.0 | 4.6 |

**Table 1B:** α for different conditions on chain alignments for randomised data

| Cycle | $\theta = \phi = 45^\circ$ | $\theta = 45^\circ, \phi = 30^\circ$ | $\theta = 30^\circ, \phi = 45^\circ$ | $\theta = \phi = 30^\circ$ |
| --- | --- | --- | --- | --- |
| 10 | 4.0 | 5.1 | 4.7 | 4.6 |
| 11 | 4.1 | 5.0 | 4.8 | 6.0 |
| 12 | 3.8 | 4.7 | 4.7 | 5.5 |
| 13 | 3.6 | 4.8 | 4.4 | 5.6 |

### Spindle alignment in explants

Extracts initially containing 2, 3 and 4 dividing nuclei were analysed in terms of spindle axis orientation by analysis of microtubule reporters at the onset of anaphase B. Using MATLAB^®^ home-made scripts, the two minor orthogonal angles (*ϕ*_1_ and *ϕ*_2_) were determined by manual clicking at spindle poles. These angles can range between 0° (parallel orientation) and 90° (perpendicular orientated).

### Dumbbell model of nuclear alignment in explants

For the model, we consider a simple phenomenological model: 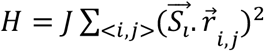, where 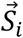 represents the orientation of aster *i* (with |*S_i_*| = 1) and 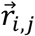 is the unit vector between aster *i* and its nearest neighbours *j*. The model is quadratic as there is no preferential direction for *S*. In this case, energy is minimised if nuclei align perpendicular to the vector of separation, 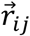, between nuclei. In the case of two nuclear spindles, it is clear this results in parallel aligned nuclear spindles, both of which are perpendicular to the vector between the nuclei. For three spindles, positioned at (0,0), (1,0) and 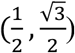, the energy is minimised for 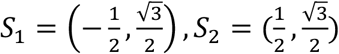 and *S*_3_ = (1,0) (direction of the vectors can also be inverted). We use a Metropolis algorithm to simulate the alignment of asters at different effective temperatures *T*. There is only one parameter, defined by *J/k_B_ T*, which represents the competition between alignment force (*J*) and random fluctuations (*k_B_T*). In Suppl. Fig. 3D–E, we show simulation outputs for 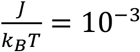 and 10°. Result presented in Fig. 3 are for 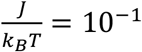

Of course, we can consider more complex models, such as 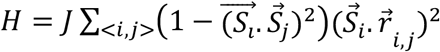 which incorporate both terms involving neighbouring aster alignment and their alignment relative to 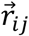. There is also similarity to models of nematic ordering in liquid crystals^34^, which have recently been applied to other biological systems^81^. Studies of self-propelled particles with repulsive interactions are also relevant, where longer ranged interactions are also considered^82^. Our aim here is to simply show how simple dumbbell-like repulsion (which results in one rotational degree of freedom) can lead to different behaviours depending on the system topology, and not to build a precise model for how such potentials interact.

### Dynamic model of aster interactions

The cytoplasm is viscous. For a viscous material, the velocity, *v*, of an object is dependent on the applied force *F*: *v* ≈ *γF*, where *γ* is the effective viscous drag coefficient. In our simple dynamic model implemented in Matlab^®^ we consider *γ = 1* and isolated asters with a circularly-symmetric force potential described by *f*(*r*) = *f_slip_*(*r*) × *ρ_MT_*(*r*), where *r* is the distance from the aster centre (centrosome), 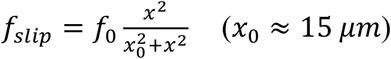 and *ρ_MT_*(*r*) represents the distribution of microtubules from the aster. We incorporate *f_slip_* to account for the reduced apparent microtubule force generation at short distances. For simplicity, we take the same characteristic distance *x*_0_ for both aster-boundary and aster-aster interaction. To account for boundary conditions, we introduce a mirror charge outside the circle for each aster.

For single asters, we only consider interactions between the wall and aster. We take *ρ_MT_*(*r*) = *e^−r/λ^*, with *λ* = 10μm and *f*_0_ = 0.01 and *r* is the perpendicular aster-wall separation. We also include a ‘noise’ term, *δf* = 0.0025. So, 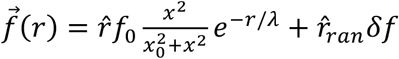 where 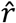 is the unit vector between aster and wall, and 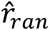 is a random unit vector generated at each time iteration. For two asters, the force is given by 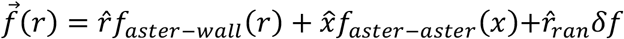. *f_aster_−wall*(*r*) is the same as for the one aster scenario. 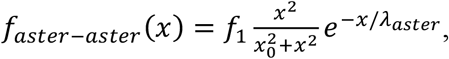 where *x* is the aster-aster separation and 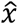 is the unit vector between the two asters, *f*_1_ = 0.0075 and *λ_aster_* = 8μm.

Considering the aster-aster separation (Suppl. Fig. 6F), we assumed the aster pair initially separated by 2 μm and centred within the *in silico* explant space. For the single aster case, we randomly initialised the aster position within the space. For Suppl. Fig. 4G–I, we initialised the aster positions randomly. Simulations were always run until the aster position reached a steady-state and angles between asters were measured at the last time point.

### Analysis of free asters in explants – distance distributions

Distance between asters and from aster to the boundary were obtained in explants at steady state, *i.e*. where asters did not move anymore (usually 30–45 min after explant deposition). The inter-aster distance was determined as Euclidean distance in 3D. We defined the *boundary distance* (*b*, *b*_1_, *b*_2_) as the shortest distance from the aster to the interface between glass, oil and cytosol, determined manually using the FIJI measurement tools (at a precision of ±0.5 μm). To determine the explant boundary on the glass (approximated with a circle of radius *R*), maximum intensity projections of both fluorescence emission channels was assessed to trace the interface between the glass, oil and cytosol. For larger explants with high aspect ratio – a quasi-2D situation – the definition of *boundary distance* served as good approximation for a boundary in two dimensions. However, in small explants where the aspect ratio is not as high, two asters sometimes aligned considerably in the third dimension. In these cases, the definition for *boundary distance* led to an underestimation of the maximum projected inter-aster distance *M* = 2*R* − *b*_1_ − *b*_2_; it becomes a geometric problem in 3D and the longest dimension is not necessarily in the plane of the glass-explant interface. This is evident for some data points in small explants (yellow dots in Fig. 4B). Finally, a correlation analysis of boundary distances *b*_1_ and *b*_2_ in the two-aster scenario (Suppl. Fig. 4F) was calculated using Pearson’s *r* in MATLAB^®^.

### Analysis of free asters in explants – dynamics

The coordinates of free asters were obtained by applying a Gaussian blur filter (radius: 1–2 pixels) and using the plugin TrackMate v3.5.1 of Fiji ImageJ^79,80^. The coordinates of detected spots were imported into MATLAB^®^ for assignment and distance calculation similarly as mentioned above.

The instant relative velocity was calculated using the formula: 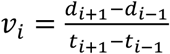, where d is the 3D Euclidian distance and t is time in the flanking time points of the measure point.

For unperturbed experiments, data was normalised to the maximum distance achieved in the separated phase in order to correct for scaling effect during splitting dynamics (Fig. 5). This data was fitted to the phenomenological equation (Suppl. Fig. 6B):

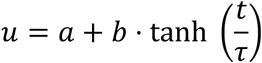

To analyse the lipid droplets, we performed a similar analysis using Fiji TrackMateJ^79,80^. Seven extracts were analysed with an aster present, with over 100 individual tracks of lipid droplets. RMS distance was then extracted across the entire time course of imaging. Similar analysis was performed in extracts without an aster. Curves (Suppl. Fig. 5H) were fitted using the in MATLAB^®^, with r^2^ = 0.98 (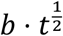 for no-aster data), r^2^ = 0.96 (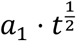 for 1-aster data), and r^2^ = 0.99 (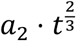 for 1-aster data). Fitting the general function *a* · *t^C^* gives a best fit for *c* = 0.66 for the one aster data.

### Microtubule profile quantification

For single asters (Fig. 5A–C), we quantified the microtubule intensity using the intensity of the RFP::β-Tubulin signal. Taking the point when asters were either 5 μm or 20 μm from the explant boundary, we used FIJI to measure the microtubule intensity along a 10 μm straight line from the edge and through the aster. The line had a width of 2 μm. For each experiment, we normalised the total intensity by the maximum measured value and then binned the data in 0.2 μm bins. Hence, the recorded intensity does not reach one, and the mean intensity only reaches a maximum around 0.8 as the maximum value does not occur at the same position.

Similar analysis was performed for the scenario with two asters (Fig. 5D–G). In this case, the centroids of the asters were used to define a straight line along which the microtubule intensity was measured throughout the process of aster separation. From this straight line between the asters, we also generated the kymograph shown in Fig. 5F right.

### Analysis of free asters in explants –perturbations with drugs and UV ablation

For comparison between control and perturbation experiments, data was time-aligned to the perturbation time-point (t=0) and plotted as average ± s.d. from at least three replicates for each condition. The change of inter-aster distance during the first 3 s after drug injection was estimated by linear regression assuming normally distributed noise, and the confidence interval of the estimated slope served as test statistic for differences between control and perturbation. Differences in final, steady-state inter-aster distance were tested by comparing the pools of distances from the last 3 s (=12 frames), using Wilcoxon signed-rank test. A significance level of 0.05 was defined prior to testing. In the case of UV ablations, the position of the aster five frames before ablation was defined as coordinate origin. The two main axes of the ellipsoid, along which the pulsed ablation was performed, defined the cartesian coordinate system. A displacement vector of the current aster position relative to the origin was calculated for each time point. The mean and standard deviation of axial (Δx) and lateral (Δy) displacement was plotted in time (Fig. 6B–C).

### Analysis of nuclei internalisation in embryos – angle probability distributions from cuts

Embryos expressing H2Av::mCherry were segmented using level sets and watershed algorithms in MATLAB^®^. Regions of low nuclear density were identified as pixels that were positioned greater than 20% of the average nucleus separation from the nearest nucleus. The centre of mass of the low-density region was identified. The division angle orientation *φ* of the neighbouring nuclei was measured relative to the centre of mass. Therefore, a nucleus dividing directly into the region of low density would be assigned an angle of 0°, and a nucleus dividing perpendicular to the region would be assigned an angle 90°. As the division does not have a preferred direction, the angle range is between 0° and 90°. A similar analysis was performed for the laser ablations, where the centre of the low-density region (artificially generated by ablating nuclei) was used to determine the relative angle of the division axis for the neighbouring nuclei.

## Acknowledgements

We thank members of the I.A.T. and T.E.S. labs for fruitful discussions, and Virgile Viasnoff, Gianluca Grenci, Tetsuya Hiraiwa, Jacques Prost and Thomas Surrey for constructive comments and feedback. We thank the staff of the Fly Facility, the Advanced Imaging Facility (AIF) and the Technical Support Service at the Instituto Gulbenkian de Ciência (IGC). Transgenic fly stocks were obtained from Bloomington Drosophila Stock Center (NIH). We acknowledge financial support from: Human Frontiers Science Program (HFSP), awarded to I.A.T. and T.E.S. and supporting J.C. and S.T. (RGY0083/2016); Fundação Calouste Gulbenkian (FCG), the Fundação para Ciência e a Tecnologia (FCT) supporting I.A.T. (Investigador FCT IF/00082/2013); EU FP7-PEOPLE-2013-CIG (N° 818743) awarded to I.A.T. and supporting J.C.; LISBOA-01-0145-FEDER-007654 supporting IGC’s core operation; LISBOA-01-7460145-FEDER-022170 (Congento) supporting the Fly Facility; PPBI-POCI-01-0145-FEDER-022122 supporting the AIF, all co-financed by FCT (Portugal) and Lisboa Regional Operational Program (Lisboa2020) under the PORTUGAL2020 Partnership Agreement (European Regional Development Fund).

## Author contributions

J.C., T.E.S. and I.A.T. conceived the study; I.A.T. and J.C. designed the experiments; J.C. carried out all experiments, and independently confirmed the laser ablation experiments in embryos, which were carried out and analysed earlier by T.E.S., jointly designed with L.H.; S.T. conceived, designed and performed the force chain analysis, assisted with image analysis, and contributed to the biophysical modelling; T.E.S. designed and performed the dumbbell model simulations, and performed the quantification of aster dynamics; S.T. and T.E.S. developed and implemented the dynamic aster repulsion model. J.C. and I.A.T. performed the quantification of all other experimental data from embryos and explants; I.A.T. designed and assembled the experimental setup; J.C., T.E.S. and I.A.T. wrote the manuscript. T.E.S. and I.A.T. jointly supervised the project.

## Data availability

All original data is available upon request.

## Code availability

All the codes developed in this study leading to the results from experiments or modelling are made available upon request.

## Supplementary Figures

**Supplementary Fig. 1 –.**
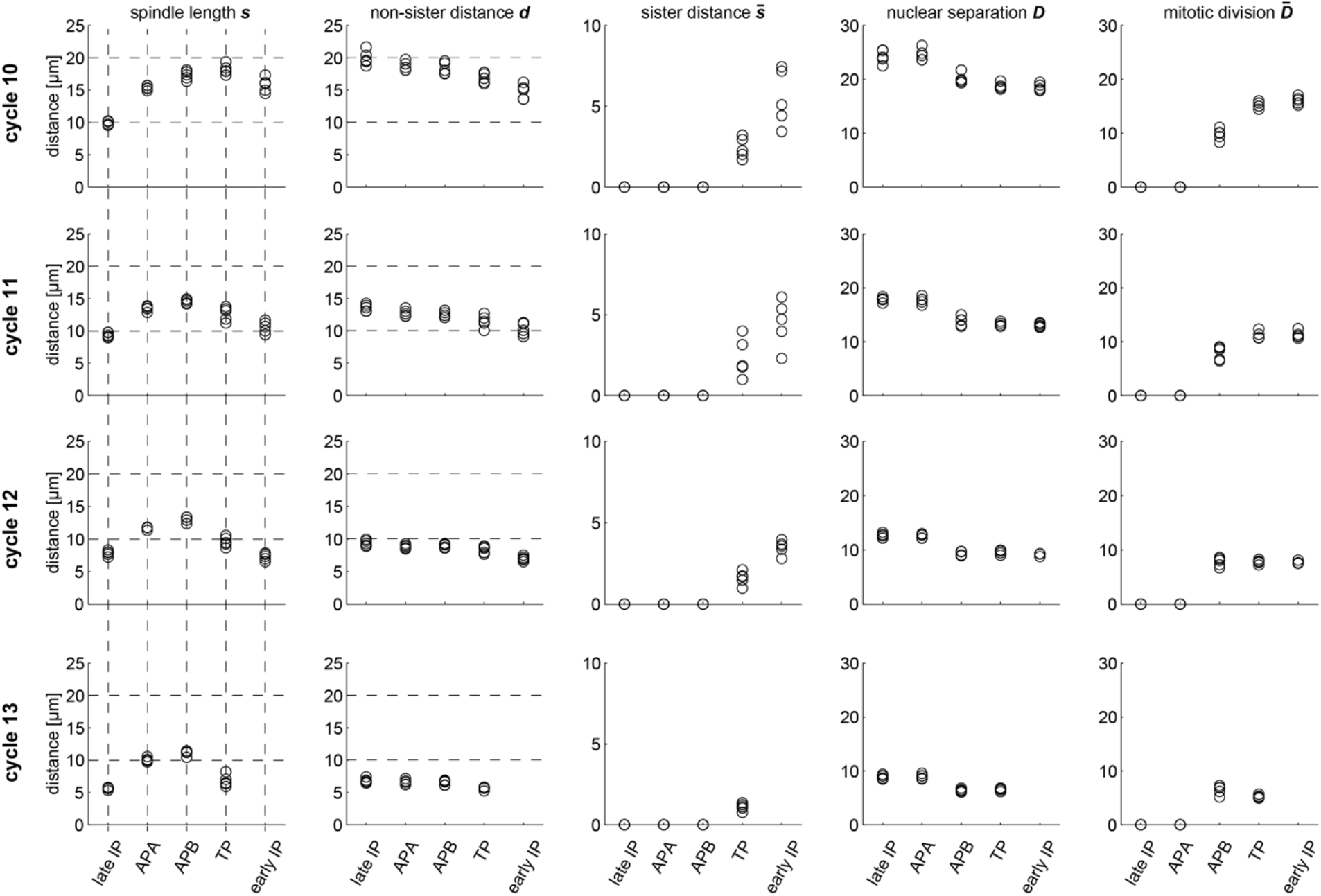
Timeline of average inter-nuclear and inter-aster distances. Relates to Fig. 1. Averages from five blastoderm embryos during cycles 10, 11, 12 and 13. Each cycle is characterised by late interphase (late IP), anaphase A (APA), anaphase B (APB), telophase (TP) and early interphase (early IP). Morphological criteria for identification of mitotic phases and hierarchical classification of distances; *Nuclear-based: D* – non-sister nuclei identified as nearest neighbours. 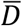 – sister chromatids/nuclei. *Aster-based: s* – sister asters ~ spindle length. 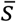 – sisters in the following mitotic cycle, i.e, *s* becomes 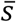 after early interphase of the ensuing mitotic cycle. *d* – non-sister asters identified as nearest neighbours. The distribution data from cycle 10 is presented in Fig. 1, and details on the measurements are described in Methods.

**Supplementary Fig. 2 –.**
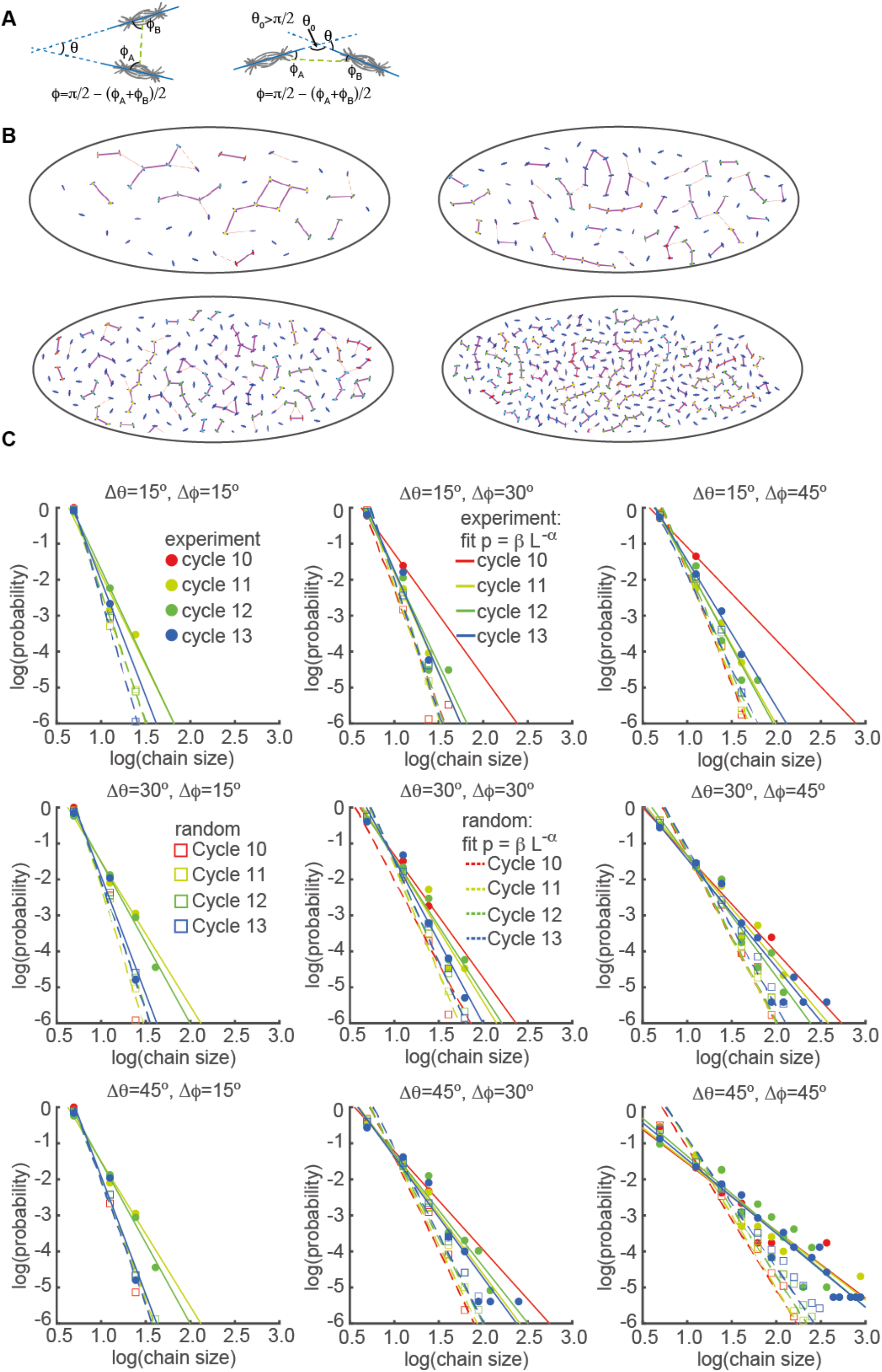
Scaling of chain size probability with chain size for different inclusion criteria. Relates to Fig. 2. **(A)** Schematic of geometric considerations in 2D of neighbour spindle alignment. Spindle axes are in cyan. Left shows almost parallel division axes (*θ* small), orientated perpendicular to the vector between the two spindles (*ϕ* almost 90°). Right shows almost colinear alignment (*θ* small, *ϕ* small). The spindle does not have a specific vector direction, so angles are between 0° and 90°. A chain is defined by spindles fulfilling one or two of the following conditions: (i) 0 ≤ *θ* ≤ Δ*θ* (ii) Δ*ϕ* ≤ *ϕ* ≤ 90° (see Methods). **(B)** Representative embryo in n.c. 10, 11, 12 and 13 showing alignment chains. Solid purple lines denote nearest neighbours that satisfy the two chain conditions with angles *Δϕ =* 45° and *Δθ =* 45°. Thin dashed red lines denote nearest neighbours that satisfy only chain condition (i). **(C)** Logarithmic plots of probability as a function of chain size (number of nuclei belonging to the chain) for different chain conditions. Filled symbols denote experimental data and open symbols are from simulations of randomised orientations. Lines represent a fit to *βL^−α^*, where *L* is the chain size and *α* is the scaling exponent (fitting performed with *fit* in Matlab^®^); solid lines are fits to experimental data and dashed lines are fits to simulated data of randomised orientations. We generated 1000 random orientations for each embryo at each cycle.

**Supplementary Fig. 3 –.**
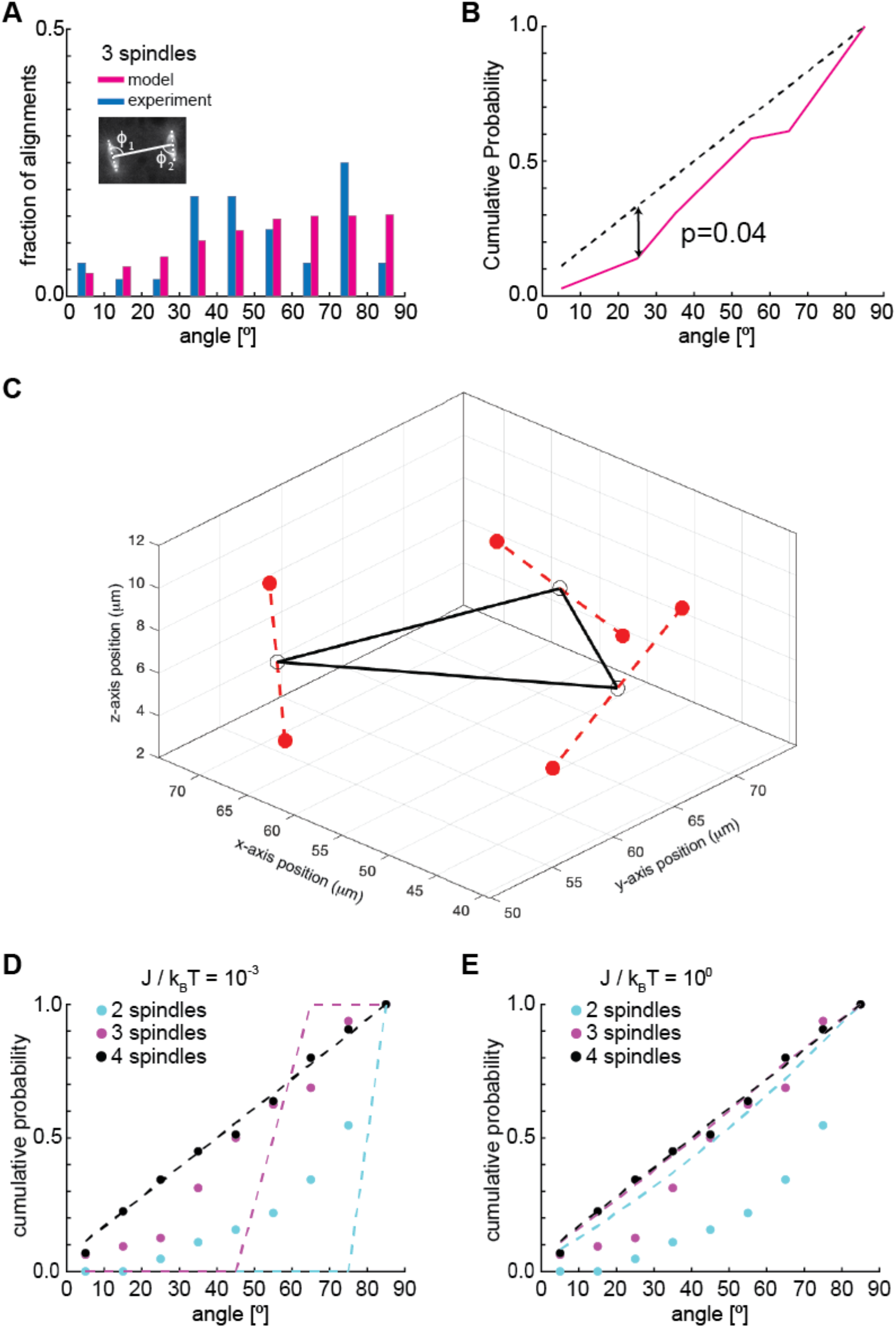
Division orientation of three spindles in a cytosolic explant, and influence of effective temperature. Relates to Fig. 3. **(A)** Histogram of the angle *ϕ* between spindle axis and the connecting line (inset) for the three-spindle scenario from experimental (blue, n=7) and *in silico* (magenta) data with stochasticity parameter J/k_B_T = 10^−1^). **(B)** Cumulative probability of relative division axis for the 3-spindle scenario and randomised orientation. p-value was determined from Kolmogorov–Smirnov test. **(C)** Three-dimensional plot of the shortest distance (black solid line) between three spindle centres (open circles). The spindle orientations (dashed lines) and positions of associated asters (filled circles) illustrate the adjustment of orientation and position in this simple system. **(D, E)** Cumulative probabilities of spindle axis orientation for two-, three- and four-spindle scenarios at different effective simulation “temperature” (stochasticity introduced as thermal noise). Dashed lines are model predictions with either low (**D**) or high (**E**) effective temperature.

**Supplementary Figure 4 –.**
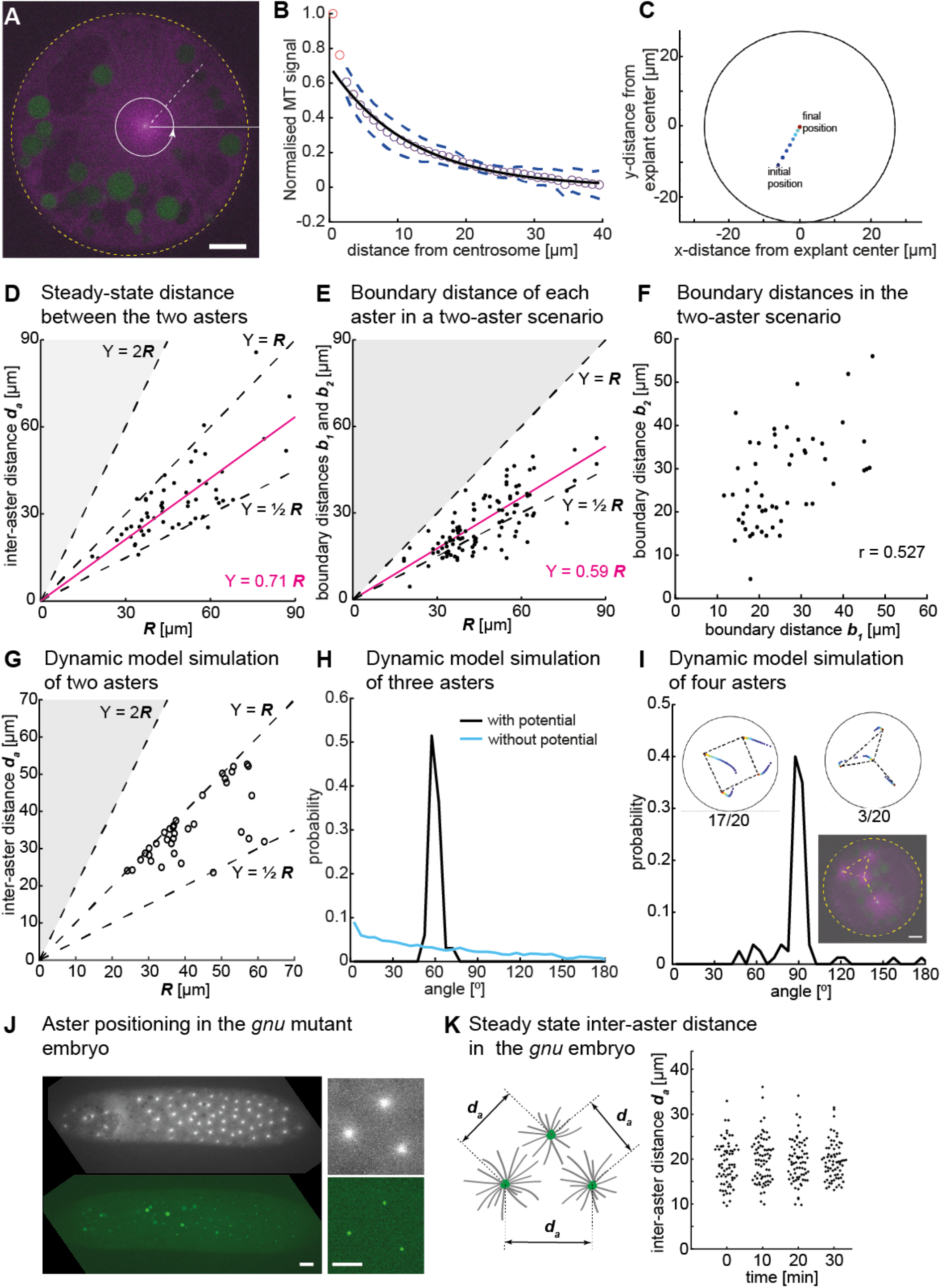
Distance analysis of asters in explants and embryos in the absence of dividing nuclei. Relates to Fig. 4. **(A)** Single Z-plane image of an explant from a *gnu* mutant embryo expressing RFP::β-Tubulin (magenta) and Spd2::GFP (green), containing a single aster. The dashed white line and the circular arrow represent the radial maximum intensity projection of the microtubule signal from the centrosome towards the periphery aiming at measuring aster size. The yellow dashed circle represents the explant boundary. Scale bar, 10 μm. **(B)** Normalized intensity of astral microtubules as schematically outlined in **A**. The black line is a mono-exponential fit to the data excluding the first two data points (red), representing the centrosome, and the dashed lines mark ±1s.d. The decay length is 11.8 ± 0.5 μm (mean ± s.e.m.), and the intensity drops to background level at ~40 μm. **(C)** Dynamic model simulation of a single aster in a circular space similar to explants in experiments. Asters always moved towards the centre. The example shows a space with *R* = 30 μm in which the final position is the centre. In larger spaces asters do not reach the centre but move only up to the interaction distance of the force potential. **(D)** Scattered plot of inter-aster distance (*d_a_*) as a function of the radius (*R*) of explants containing two asters (n=54). Most measured data points fall between the dashed lines denoting the explant radius (*Y = R*) and half of the radius 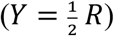. The magenta line represents the linear regression. **(E)** Scatter plot of shortest distance to explant boundary (*b*_1_ and *b*_2_) as a function of the radius *R* in explants containing two asters (n=54; details in inset of Fig. 4B). The magenta line represents the linear regression. Black dashed lines represent half and full radius distance. **(F)** Correlation plot of the boundary distances *b*_1_ and *b*_2_ referred to in panel E with Pearson’s correlation coefficient *r*. **(G)** Dynamic model simulation of the 2-aster scenario in *in silico* explants with varying size *R*. Simulations are in agreement with experiments shown in **D**. **(H)** Angle distribution from aster positions in a dynamic model simulation with three asters. The simulation evolved from initially random positions, and asters robustly moved towards a triangular configuration, as shown in Fig 4C. The peak at 60° represents equal distances between the three asters (Methods). In the absence of a repulsion potential the regularity is lost (blue line). **(I)** Angle distribution from aster positions in a dynamic model simulation with four asters. The two insets show the temporal evolution of position (color-coded as **C**) and the final configuration marked with dashed lines. The majority of simulations (17/20) resulted in a regular square (top left inset) with 3/20 resulting in a “Y” configuration (top right inset). These configurations were also observed in embryo explants (Fig. 4C and bottom inset). **(J)** Maximum intensity Z-projection of a *gnu* mutant embryo expressing RFP::β-Tubulin (grey) and Spd2::GFP (green) (left, scale bar 20 μm), with magnification of three asters (right, scale bar 10 μm). **(K)** Schematic of the measurement of inter-aster distance *d* between nearest neighbour asters (left), and scatter plots of *d_a_* during consecutive intervals of 10 min for the same embryo.

**Supplementary Figure 5 –.**
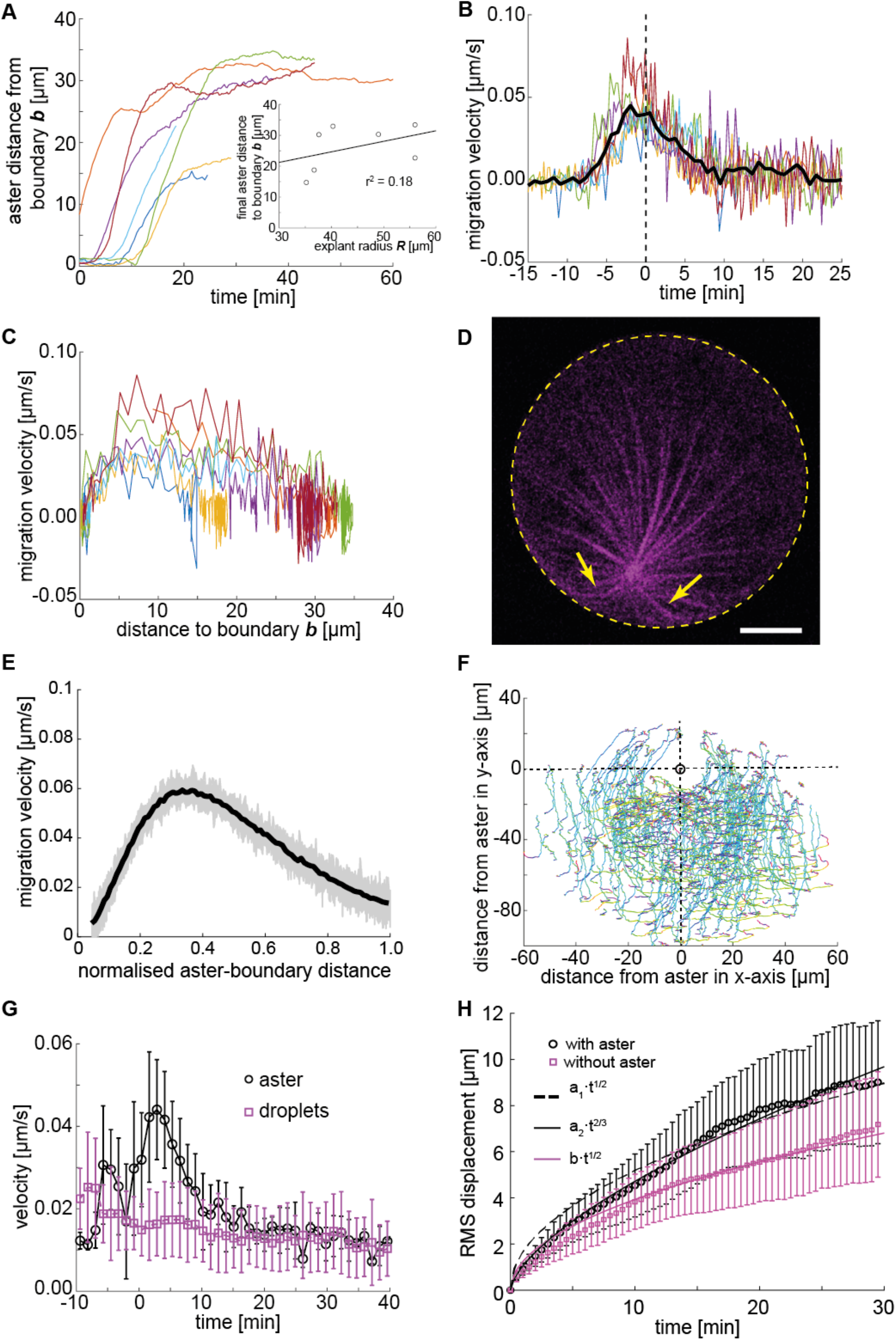
Analysis of migration dynamics in explants containing a single aster. Relates to Fig. 5. **(A)** Trajectories of aster distance to the explant boundary from independent experiments. The inset shows correlation between explant radius *R* and final aster distance to the explant boundary after reaching a steady state. The solid line represents the linear regression to experimental data with Pearson’s correlation coefficient *r*. **(B)** Migration velocity as a function of time, where t=0 is defined as the time when the aster lies midway between the explant edge and the final position of the aster. Solid line represents average over all measurements. **(C)** Migration velocity as a function of distance to the boundary. **(D)** Z-projection of a 3D image stack of a small explant containing one aster that exemplifies microtubule buckling and splay near the explant boundary represented by the yellow dashed circle (scale bar 5 μm). **(E)** Migration velocity as a function of normalised boundary distance obtained from a dynamic model simulation; individual velocity profiles (n=100, grey) and average (black) are shown, in good agreement with experimental data (Fig 5B). **(F)** Velocity field of yolk droplets around the single aster as it escapes from the explant boundary. Image rotation and frame matching were performed to overlay all experiments such that the aster, at each time point is positioned at (0,0) and moves in the direction (0,-1). Track colour coding denotes displacement angle, where turquoise corresponds to 0° and red to 90°. **(G)** Velocity profile of asters (black) and 35 lipid droplet (magenta) tracks within a distance of 20 μm from the aster, where t=0 is the time point at which the aster starts moving away from the explant boundary. **(H)** Root-mean-square (RMS) displacement of lipid droplets in the explants. Droplet movement analysed with (black) and without (magenta, n=6) an aster. Error bars s.e.m. and lines represent fits to models shown in legend. Measurements in explants with aster fit best to a model exhibiting some directionality, see Methods for further discussion.

**Supplementary Figure 6 –.**
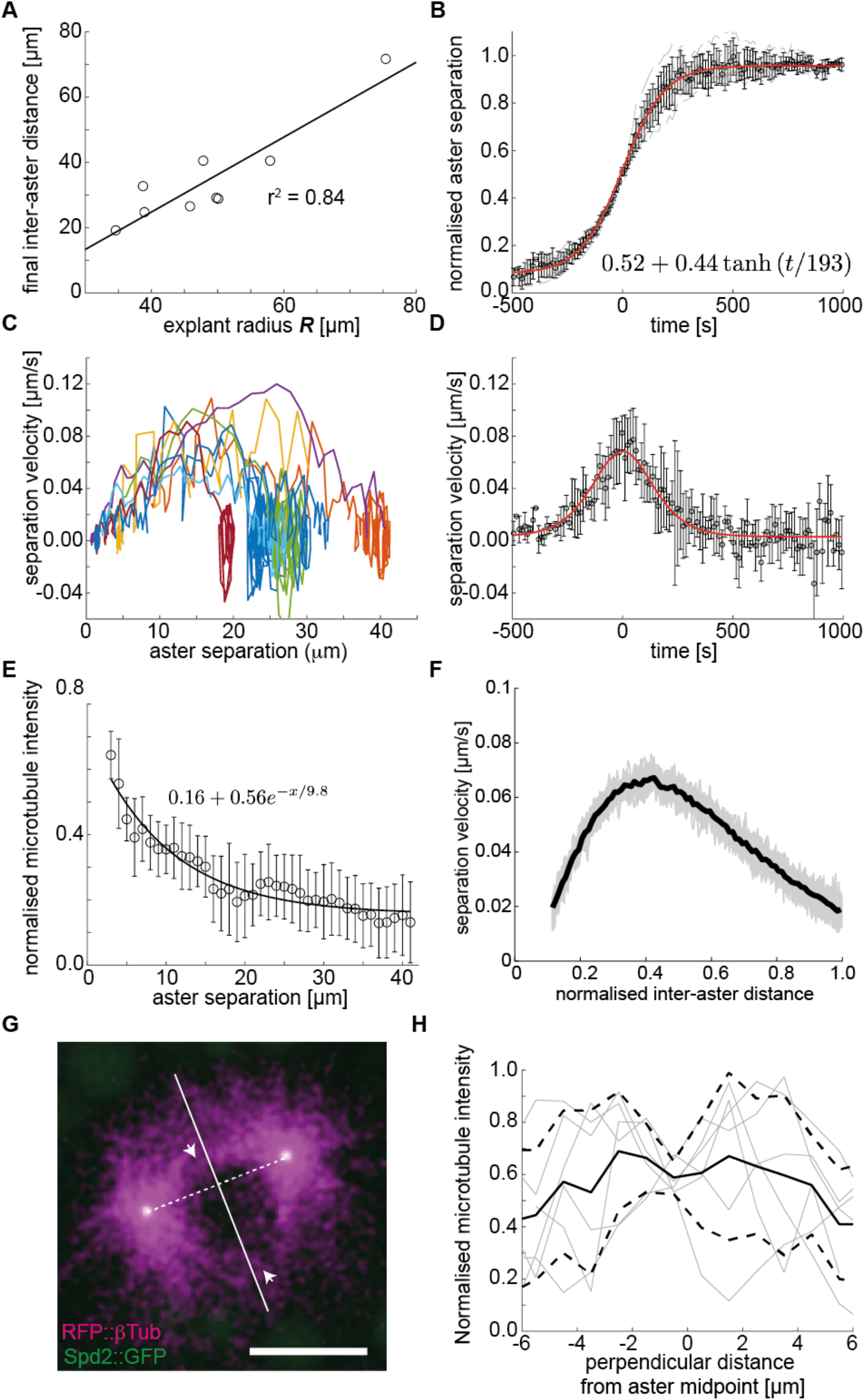
Analysis of separation dynamics in explants containing two asters. Relates to Fig. 5. **(A)** Correlation between explant radius *R* and the final distance between asters after reaching a steady state (n=9). The solid line represents the linear regression to experimental data. **(B)** Normalised aster separation distance versus time, where t=0 is defined as the time when the aster lies midway between the explant edge and the final position of the aster. Solid line represents the fit to a hyperbolic tangent function (n=9). **(C)** Aster separation velocity as a function of aster separation, as defined above. Each colour corresponds to a different experiment (n=8). **(D)** Aster separation velocity versus time, where t=0 is defined as above. Solid line denotes the fitting to the time derivative of the *tanh* function shown in **B** (n=9). **(E)** Normalised microtubule intensity at the midpoint perpendicular axis between asters in function of aster separation distance. Open markers denote average values and error bars the standard deviation. Solid line represents the fitting to exponential decay (n=7), fitting performed using MATLAB *fit* function. **(F)** Migration velocity as a function of normalised inter-aster distance obtained from a dynamic model simulation that does not include slippage; individual velocity profiles (grey) and average (black) are shown, in good agreement with experimental data (Fig 5E, Methods). **(G)** Two colour maximum intensity Z-projection of two separating asters in an explant with fluorescent reporters for Spd2::GFP (green) and RFP::β-Tubulin (magenta). A void of microtubules in the centre between two separating asters is observed during the initial acceleration phase. Scale bar, 10 μm. **(H)** Normalised microtubule intensity along the perpendicular midpoint axis of separating aster during the initial acceleration phase. Solid line denotes average normalised (for each experiment, as discussed in Methods) microtubule density and dashed lines ±1s.d. (n=6). There was significant variation between samples and the microtubule signal was often weak, making a detailed analysis challenging.

**Supplementary Figure 7 –.**
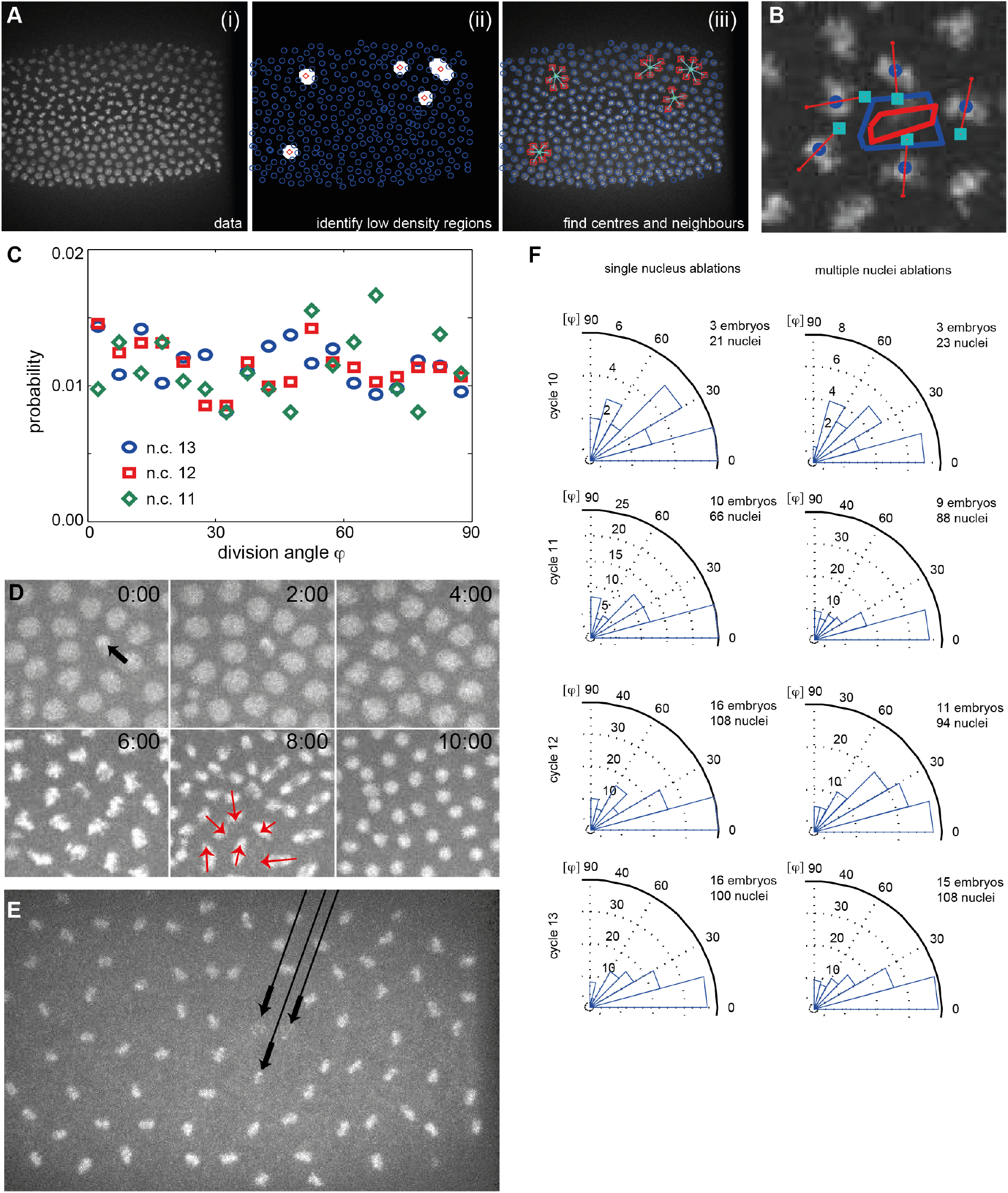
Analysis of nuclei division orientation *in vivo*. Relates to Fig. 7. **(A)** (i) Embryo expressing H2Av::mCherry just prior to anaphase B of n.c. 13 . (ii) Areas of low nuclear density were identified by finding pixels with no nuclei within 20% of the average nuclear separation and the centroid of these areas found (red diamonds). (iii) The nuclei (red squares) neighbouring the low-density region were then identified. **(B)** Examples of nuclear division near a region of low nuclear density. The division axis orientation of the neighbouring nuclei to a region of low density was measured (red bars) and the subsequent position of the nuclei after division identified (turquoise squares). Change in area of low nuclear density region from before (blue) to after (red) area is shown. **(C)** Analysis performed as in Fig. 7B, but for arbitrary locations selected within the embryo. **(D)** Artificial generation of a region of low nuclear density by ablating a nucleus (Methods). Black arrow in top left panel identifies ablated nucleus; ablation at time t=0 min. Images from maximum intensity projection of embryo expressing H2Av::mCherry. Red arrows at 8 min denote division axis orientation of neighbouring nuclei to the ablated nucleus. **(E)** Similar to **D**, except three nuclei are ablated to generate a larger region of low nuclear density. **(F)** Rose plots of division axis orientation for nuclei adjacent to ablated nuclei, for single (left column) and multi-nuclei (right column) ablations.

## Supplementary Video Legends

**Supp. Video 1:** Maximum intensity Z-projection from a 3D time-lapse movies of three distinct cycling explants starting with a single spindle extracted from embryos expressing Jupiter::GFP (grey) and H2Av: :RFP (magenta). Time in min:sec, scale bar 10 μm. Frame rate is 2 frames/min. In support of Fig. 3.

**Supp. Video 2:** Maximum intensity Z-projection from a 3D time-lapse movie of a *gnu* mutant embryo expressing β-Tubulin::EGFP (grey). The first part shows an approximately 1h old embryo, the second part an approximately 4h old embryo. Time in hr:min:sec, scale bar 20 μm. Frame rate is 2 frames/min. In support of Fig. 4.

**Supp. Video 3:** Maximum intensity Z-projection from a 3D time-lapse movie of explants generated from *gnu* mutant embryos expressing RFP::β-Tubulin (magenta) and Spd2::GFP (green). The left explant contains a single aster moving away from the explant boundary, the right explant contains two separating asters. Time in min:sec, scale bar 10 μm. Frame rate is 4 frames/min. In support of Fig. 5.

**Supp. Video 4:** Maximum intensity Z-projection from a 3D time-lapse movie of explants generated from a *gnu* mutant embryo expressing RFP::β-Tubulin (magenta) and Spd2::GFP (green), containing two separating asters, after pulse injection of solutions: control with buffer (left), 10 mM sodium azide (centre) and 0.2 mM of colchicine. Time in min:sec, scale bar 10 μm. Frame rate is 4 frames/min. In support of Fig. 5.

**Supp. Video 5:** Maximum intensity Z-projection from a 3D time-lapse movie of explants generated from a *gnu* mutant embryo expressing RFP::β-Tubulin (magenta) and Spd2::GFP (green) containing a single aster. The aster was allowed to equilibrate followed by an asymmetric elliptic ablation (yellow line at times 00:15 to 01:00) performed in control explants (no injection) and in explants supplemented with 0.2mM of colchicine. Time in min:sec, scale bar 10 μm. Frame rate is 4 frames/min. In support of Fig. 6.

**Supp. Video 6:** Maximum intensity Z-projection from a 3D time-lapse movie of an explant containing two separating asters from a *gnu* mutant embryo expressing RFP::β-Tubulin (magenta) and Spd2::GFP (green). The elliptic ablation (yellow line from 00:15 to 00:45) was performed when asters were ~7 μm apart. Time in min:sec, scale bar 10 μm. Frame rate is 4 frames/min. In support of Fig. 6.

**Supp. Video 7:** Maximum intensity Z-projection from a 3D time-lapse movie of a wildtype embryo expressing H2Av::mCherry (magenta) and Spd2::GFP (green) during n.c. 12 and 13, in response to spontaneous nuclear internalisation. Time in min:sec, scale bar 20 μm. Frame rate is 2 frames/min. In support of Fig. 7.

